# *Zymoseptoria tritici* stealth infection is facilitated by stage-specific down-regulation of a β-glucanase

**DOI:** 10.1101/2024.12.04.626787

**Authors:** Diego Rebaque, Cristian Carrasco-López, Parvathy Krishnan, Gemma López, Sergio López-Cobos, Felipe de Salas, Lukas Meile, Cécile Lorrain, Asier Largo-Gosens, Bruce A. McDonald, Francisco Vilaplana, Maria Jesus Martinez, Hugo Mélida, Antonio Molina, Andrea Sánchez-Vallet

**Affiliations:** Centro de Biotecnología y Genómica de Plantas, Universidad Politécnica de Madrid (UPM) - Instituto Nacional de Investigación y Tecnología Agraria y Alimentaria (INIA/CSIC), Campus de Montegancedo UPM, Pozuelo de Alarcón (Madrid), Spain; Division of Glycoscience, School of Engineering Sciences in Chemistry, Department of Chemistry, KTH Royal Institute of Technology, Stockholm, Sweden; Plant Pathology, Institute of Integrative Biology, ETH Zürich, Zürich, Switzerland; Área de Fisiología Vegetal, Departamento de Ingeniería y Ciencias Agrarias, Universidad de León, León, Spain; Centro de Investigaciones Biológicas Margarita Salas, Spanish National Research Council, C/ Ramiro de Maeztu 9, 28040 Madrid, Spain; Instituto de Biología Molecular, Genómica y Proteómica (INBIOMIC), Universidad de León, León, Spain; Departamento de Biotecnología-Biología Vegetal, Escuela Técnica Superior de Ingeniería Agronómica, Alimentaría y de Biosistemas, UPM, Madrid, Spain

**Keywords:** Plant cell wall, pathogen effector, plant resistance, effector gene regulation, Damage-associated molecular pattern (DAMP)

## Abstract

Plant cell walls constitute a major defence barrier against pathogens, although it is unclear how specific cell wall components impact pathogen colonisation. Pathogens secrete cell wall degrading enzymes (CWDEs) to facilitate plant colonisation, but damaged, infected cells are often a source of cell wall-derived oligosaccharides that trigger host immunity. The mechanisms by which pathogens minimize the release of cell wall-derived oligosaccharides while colonizing the host remain to be elucidated. We combined biochemical, molecular genetics and transcriptomic analyses to functionally characterize a glycoside hydrolase (*Zt*GH45) from the wheat pathogen *Zymoseptoria tritici*. *ZtGH45* gene is expressed during the necrotrophic phase of the fungus, coinciding with an accumulation of wheat β-1,3/1,4-mixed-linked glucan (MLG)-derived oligosaccharides. We show that overexpression of *ZtGH45* enhances β-1,3/1,4-glucan hydrolysis and the derived oligosaccharides trigger an immune response in wheat, which hinders *Z. tritici* virulence. The results demonstrate that tight regulation of *ZtGH45* is critical for the infection process to prevent early accumulation of MLG oligosaccharides that would prematurely induce host immunity counterbalancing fungal virulence. We suggest that the balance between plant cell wall degradation by fungal CWDE and the release of immunogenic wall-derived oligosaccharides governs the outcome of host invasion by pathogens.

## Introduction

Successful plant colonisation largely depends on the pathogen’s ability to overcome the host resistance machinery and efficiently acquire nutrients (Haueisen & Stukenbrock, 2016). The plant cell wall acts as a defensive barrier against potential pathogen attacks but also represents a source of nutrients and biosynthetic building blocks for colonising microorganisms. To surmount this defensive barrier and acquire nutrients, fungal pathogens secrete an arsenal of cell wall degrading enzymes (CWDEs). CWDEs are a subset of carbohydrate active enzymes (CAZymes, Drula et al., 2022), including glycosyl hydrolases (GHs), which hydrolyse the glycosidic linkage of carbohydrates present in the plant cell walls. The activity of CAZymes can lead to the release of plant cell wall-derived glycans, which can function as elicitors of host immune responses and are commonly known as damage-associated molecular patterns (DAMPs; Molina et al., 2024a).

Plant cell wall-derived glycans can be perceived by plant Pattern Recognition Receptors (PRRs), triggering a cascade of defence responses referred to as pattern triggered immunity (PTI) that include the accumulation of reactive oxygen species (ROS), stomatal closure, callose accumulation and up-regulation of defensive genes (Bigeard *et al*., 2015; Boutrot & Zipfel, 2017; Gust *et al*., 2017; Bacete *et al*., 2018; Molina *et al*., 2024a). Thoroughly investigated plant cell wall-derived oligosaccharides acting as DAMPs include oligogalacturonides (OGs) derived from pectins (Hahn *et al*., 1981; Voxeur *et al*., 2019), as well as oligosaccharides derived from xyloglucans (Claverie *et al*., 2018), mannans (Zang *et al*., 2019), callose (Mélida *et al*., 2018), arabinoxylan (Mélida *et al*., 2020), xylans (Pring *et al*., 2023; Fernández-Calvo *et al*., 2024), cellulose (Aziz *et al*., 2007; Martín-Dacal *et al*., 2023) and β-1,3/1,4-glucan (Rebaque *et al*., 2021; Yang *et al*., 2021). β-1,3/1,4-glucan, also known as mixed-linked glucan (MLG), consists of linear polymers of glucose molecules linked by β-1,4 and β-1,3 bonds, and is a distinctive feature of the plant cell wall of Equisetaceae and Poaceae species (Labavitch & Ray, 1978; Sørensen *et al*., 2008; Fry *et al*., 2008). MLG oligosaccharides can also be derived from walls of oomycetes and be perceived by the host as microbe-associated molecular patterns (MAMPs) (Rebaque *et al*., 2021). Regardless of their origin, MLG-derived oligosaccharides, particularly the well- characterised trisaccharide β-ᴅ-cellobiosyl-1,3-β-ᴅ-glucose (MLG43), confer enhanced disease resistance to pathogens in several plant species, including Arabidopsis, tomato, pepper, barley, and rice (Rebaque *et al*., 2021; Barghahn *et al*., 2021; Yang *et al*., 2021). Recent works have shown that the LysM receptor kinase CERK1 (Chitin Elicitor Receptor Kinase 1), and Lectin Receptor Kinase 1 (LecRK1) are involved in the perception of MLG oligosaccharides in rice (Yang *et al*., 2021; Dai *et al*., 2023) and that the Leucine Rich Repeat-Malectin Receptor Kinases (LRR-MAL RKs) IGP1/CORK1, IGP3 and IGP4, and the LysM-containing receptors CERK1, LYK4 and LYK5 are required for perception of MLG oligosaccharides in Arabidopsis (Rebaque *et al*., 2021; Martín-Dacal *et al*., 2023).

Septoria tritici blotch is one of the major diseases of wheat globally and is caused by the fungal pathogen *Zymoseptoria tritici.* This fungus harbours in its genome around 200 genes encoding putative glycoside hydrolases (GHs), which potentially function as CWDEs (Drula *et al*., 2022). The initiation of the infection cycle in *Z. tritici* occurs upon the germination of either a sexual or an asexual spore on the surface of plant leaves. Following stomatal penetration, the fungus colonises the apoplastic space in intimate interaction with plant cell walls without exhibiting any visible signs of plant damage for a period ranging from 8 to 20 days post infection (dpi; (Kema *et al*., 1996; Duncan & Howard, 2000; Keon *et al*., 2007; Deller *et al*., 2011; Sánchez-Vallet *et al*., 2015; Steinberg, 2015). Effector recognition by wheat resistant cultivars prevents *Z. tritici* penetration through the stomata and the subsequent apoplast colonization (Battache *et al*., 2022; Alassimone *et al*., 2024) Coincident with the onset of necrotic symptoms, fungal growth spikes and the reproductive structures known as pycnidia and pycnidiospores are produced (Steinberg, 2015). Distinct sets of virulence factors are postulated to be required throughout the different phases of a *Z. tritici* infection. In fact, several predicted cellulases, hemicellulases, pectinases, and cutinases show a specific life-cycle-dependent expression pattern during wheat infection (Brunner *et al*., 2013; Palma-Guerrero *et al*., 2017). However, the mechanistic contribution of CWDEs to *Z. tritici* virulence remains largely unknown. In other pathogens, several CWDEs have been shown to play key roles in the infection process. For example, impairment of the cellulose degradation machinery in the soil-borne fungus *Fusarium oxysporum* increases its virulence in Arabidopsis (Gámez-Arjona *et al*., 2022). Similarly, it was shown that constitutive overexpression of two endoglucanases from the GH12 family (MoCel12A and MoCel12B) in the rice pathogen *Magnaporthe oryzae* results in an increased release of MLG oligosaccharides and a reduction of symptom development (Yang *et al*., 2021). Remarkably, the contribution of CWDEs to virulence is not conserved across pathogenic fungi, though they are considered to contribute to host compatibility (Esquerré-Tugayé *et al*., 2000).

In this study, we identified a glycoside hydrolase belonging to the GH45 family from *Z. tritici* (*ZtGH45*) and demonstrated that it is tightly regulated and only expressed in the necrotrophic phase of the fungus. Overexpression of *ZtGH45* leads to enhanced accumulation of MLG-derived oligosaccharides in wheat plants infected with this fungus. These MLG oligosaccharides trigger disease resistance responses in wheat, that involve ROS accumulation and stomatal closure, which hinder *Z. tritici* infection progression. We propose that tuning the expression of CWDEs, such as *ZtGH45,* impacts pathogen colonisation and host recognition.

## Material and Methods

### Growth conditions for *Zymoseptoria tritici* and bacterial strains

The Swiss *Z. tritici* strain ST99CH_3D7 (abbreviated as 3D7; Linde *et al*., 2007) was used in this study. *Z. tritici* was grown in yeast sucrose broth (YSB, 1% w/v yeast extract and 1% w/v sucrose) or in yeast peptone dextrose (YPD, 1% w/v yeast extract, 2% w/v peptone, and 2% w/v dextrose) amended with 50 μg/mL kanamycin sulphate, at 18° C, 120 rpm in 100-mL Erlenmeyer flasks. Blastospores from *Z*. *tritici* were collected after 6 days and used for the infection and developmental assays. Liquid fungal cultures were filtered through double-layered sterile cheesecloth, blastospores were collected by centrifugation at 3273 *g* for 15 min at 4 °C and resuspended in water. Blastospore concentration in the suspension was determined using Neubauer counting chambers. For molecular cloning and plasmid propagation *Escherichia coli* strain NEB® 5-alpha (New England Biolabs) and Stellar™ Competent Cells (Takara Bio Group) were used. *E. coli* was grown on LB media (1.6% w/v tryptone, 1% w/v yeast extract, 0.5% w/v NaCl) amended with kanamycin sulphate (50 μg/mL) at 37 °C. *Agrobacterium tumefaciens* strain AGL1 was grown at 28 °C in LB media containing kanamycin sulphate (50 μg/mL), carbenicillin (100 μg/mL) and rifampicin (50 μg/mL), and used for *Agrobacterium*-mediated transformation of *Z. tritici*.

### Generation of *Z. tritici* transformants

Based on RNAseq reads (NCBI accessions: SRA SRP152081 and SRP077418), we amended the annotation of the *ZtGH45* gene in the 3D7 and 1A5 genomes. For the strains 3D1 and 1E4, the gene was correctly annotated. We obtained two types of overexpression lines, one in which the gene was inserted *in locus* and the second one with the ectopic insertion. To obtain *Z. tritici* strains overexpressing *ZtGH45* (*ZT3D7_G10118*, *ZtIPO323_110030*, *Zt09_chr_10_00532*, *Mycgr3_76589*) (Supporting Information Table S1) *in locus*, the *Aspergillus nidulans* glyceraldehyde-3-phosphate dehydrogenase (*gpdA*) promoter was inserted upstream of the start codon of *ZtGH45* by homologous recombination in the strain 3D7. Up- and down-flanking regions (approximately 1000 bp) were amplified from 3D7 genomic DNA using PfuTurbo® Cx Hotstart DNA polymerase (Stratagene, Cedar Creek, TX, US). USER Friendly cloning technique was used to fuse the flanking regions to the binary vector backbone of prfHU2E (Frandsen *et al*., 2008). Fusion was performed using USER enzyme mix from New England Biolabs (Ipswich, MA, USA) and the plasmid was cloned in *E. coli*, following the manufacturer’s instructions. To obtain *Z. tritici* strains ectopically overexpressing the wild-type or mutated version of *ZtGH45*, the plasmids were assembled using the In-Fusion HD Cloning Kit (Takara Bio, Japan). Previously, the pCCL1 plasmid was constructed by inserting the *gpdA* promoter upstream of the *Ttef* terminator into the pLM1 plasmid (Supporting Information Fig. S1; (Meile *et al*., 2020). The wild-type PCR product of *ZtGH45* (from start to stop codons) was obtained from 3D7 genomic DNA and the catalytically dead version (D37A and D145A) was constructed by PCR-driven mutagenesis (Heckman & Pease, 2007) using Supreme NZYProof 2X Green Master Mix (NZYTech). The PCR products were inserted between the *gpdA* promoter and the *Ttef* terminator of the EcoRV-linearized pCCL1 plasmid. The primers used are listed in Supporting Information Table S2. The *Z. tritici* overexpressing mutants were obtained by *A. tumefaciens*- mediated transformation and selected using hygromycin (100 µg/mL) as previously described (Zwiers & De Waard, 2001; Krishnan *et al*., 2018; Meile *et al*., 2020).

### *Z. tritici* infection assays

Wheat (*Triticum aestivum*) cultivars Titlis and Drifter (DSP Ltd., Delley, Switzerland) were grown in a growth chamber at 18 °C during the day and 15 °C during the night and 60% relative humidity for 15 days, as described (Suarez-Fernandez *et al*., 2023). A suspension of 10^7^ spores/mL of *Z. tritici* in water with 0.1% v/v Tween 20 was used to spray-inoculate wheat leaves (1 mL/plant). Upon infection, plants were placed in sealed plastic bags for 48 h to maintain high humidity. For the protection assay, 24 h before infection, wheat plants were sprayed (1 mL/plant) with a solution of 0.5 mM of β-ᴅ-cellobiosyl-1,3-β-ᴅ-glucose (MLG43; O- BGTRIB; Megazyme) or 0.5 mM cellotriose (O-CTR; Megazyme) with the adjuvants UEP-100 (0.1% v/v; Croda) and Tween 20 (0.01% v/v). The adjuvant solution alone was used as the mock control. Disease symptoms in second leaves were estimated at 13-18 dpi, as described by Meile *et al*. (2018). Leaves were analysed using ImageJ (Schneider *et al*., 2012) and an automatic image analysis method (Stewart *et al*., 2016) to estimate the percentage of leaf area covered by lesions and pycnidia density (pycnidia/cm^2^ of leaf or pycnidia/cm^2^ of lesion).

### *Z. tritici* developmental assay

A 3-µL drop at 10^6^ spores/mL of each *Z. tritici* line was placed on solid YMS (0.4% w/v yeast extract, 0.4% w/v malt extract, 0.4% w/v sucrose, and 1.2% w/v Bacto^TM^ Agar) amended with 1 mM H2O2, 1 M sorbitol, 0.5 M NaCl or 200 ng/µL Calcofluor white, or on Vogeĺs minimal media (Vogel, 1956) amended with 5 g/L sucrose, 5 g/L fructose, 5 g/L carboxymethylcellulose sodium salt (CMC; C4888; Sigma-Aldrich), or 5 g/L barley β- glucan (P-BGBL; Megazyme). Plates were incubated at 18 °C, except for one of YMS agar that was incubated at 28 °C for 6 days.

### *In vitro* assays of β-glucan enzymatic degradation

*Z. tritici* and wheat alcohol insoluble residue (AIR) was obtained as described by Rebaque *et al*., (2023) with modifications. *Z. tritici* blastospores were grown in YPD media for 6 days as indicated above. After centrifugation at 3273 *g* for 15 min at 4 °C, the cells were washed twice with distilled water. *Z. tritici* cells and dry leaves of 17 day-old wheat plants were fine-powdered and washed three times with 30 mL/g methanol/chloroform (1:1; v/v) shaking at 4 °C for 1 hour, overnight, and 1 hour. Insoluble residues were discarded after centrifuging at 3273 *g* for 10 min at 4 °C after each step. Pellets were washed with acetone for 1 hour and the resulting pellet was dried. Pellets were treated with 30 mL/g 70% v/v ethanol twice overnight and for 1 hour. Pellets were washed with acetone and the resulting dry pellet was considered AIR. For β-1,3-1,4-glucan quantification in wheat and *Z. tritici* AIR β-Glucan Assay Kit (Mixed Linkage; K-BGLU, Megazyme) was used.

To determine β-glucan oligosaccharides released by the wild-type and *ZtGH45_OE*, a spore suspension of each line of *Z. tritici* at a concentration of 4·10^5^ spores/mL was grown in Vogel’s minimal medium (Vogel, 1956) with 0.5% (w/v) fructose as carbon source to avoid presence of free glucose in the media. For the β- 1,3/1,4-glucan, cellulose and AIR degradation assay, commercial barley β-glucan (P-BGBL; Megazyme), carboxymethylcellulose sodium salt (CMC; C4888, Sigma-Aldrich) or wheat AIR were added to the media at a concentration of 0.5% (w/v). Cellulase (EC 3.2.1.4; endo-1,4-β-D-glucanase from *Aspergillus niger*; E-CELAN; Megazyme) at 2.5 U/mL was used as a positive control. The released oligosaccharides were partially purified from the supernatant from 96-hour old cultures by adding one volume of 100% ethanol and precipitating the polymers at -20 ℃ overnight (Voxeur *et al*., 2019). Samples were centrifuged at 5000 *g* for 10 min at 4 ℃ and the supernatants were collected and freeze-dried. β-glucan oligosaccharides released in the media of four different biological replicates were quantified using the modified β-Glucan Assay Kit (Mixed Linkage; K-BGLU, Megazyme). Samples were directly treated with β-glucosidase following the protocol of the manufacturer. Serial glucose dilutions were used to perform the standard curve calibration. The oligosaccharides released by the wild-type and *Zt*GH45_OE lines were compared. In addition, to identify the released oligosaccharides, samples from the commercial barley β-glucan were analysed with HPAEC-PAD in a LC 930 Compact IC Flex (Metrohm) chromatography system with an IC pulsed amperometric detector (FlexiPAD). Oligosaccharides were separated using a Metrosep Carb 2 250/4.0 (Metrohm) analytical column and a Metrosep Carb 2 Guard/4.0 (Metrohm) guard column at 40 ℃ with an isocratic 200 mM NaOH and 150 mM sodium acetate and variable flux: 0-12 min at 0,7 mL/min, 12,1-22,0 min at 0,9 mL/min and 22,1-27,0 min at 0,7 mL/min. For oligosaccharide quantification, standard curves of D-glucose (PanReac), D-fructose (Merck), MLG43 (O- BGTRIB; Megazyme), and cellotriose (O-CTR; Megazyme) commercial standards were used. The experiment was performed 2 times independently.

### Characterization of *in planta* β-glucan oligosaccharide production

For the *in planta* identification and quantification of oligosaccharide production during the course of the infection, wheat plants were infected as described above using 3D7. Two cm from the second leaf tip were discarded and the adjacent 13 cm from each leaf were collected and frozen in liquid nitrogen. Three independent biological replicates, each consisting of two second leaves per treatment, were collected for oligosaccharide analysis at 6, 8, and 10 dpi. Samples were homogenized in liquid nitrogen and freeze-dried. 10-15 mg of dry material was resuspended in 0.5 mL deionized water, ultrasonicated for 10 min and heated at 95 °C for 10 min. After cooldown, samples were centrifuged for 10 min at 17000 *g*. Supernatants were filtered through 10 KDa MWCO Centrifugal Filters (Amicon®, Ultracel®). Pellets were extracted again following the same procedure and supernatants were pooled together. Oligosaccharides-containing filtered- supernatants were freeze-dried and resuspended in 0.1% formic acid-acetonitrile (1:1 v/v). For HILIC-ESI-MS, samples were injected onto a XBridge Amide column (3.5 µm, 2.1 x 150 mm; Waters, MA, USA) with a flow of 0.8 mL/min. The mobile phase consisted of 0.1% (v/v) formic acid in water (Eluent A) and 0.1% (v/v) formic acid in acetonitrile (Eluent B). The eluent program was as follows: 20% A at 0 min, 35% A at 10 min, 55% A at 11 min, 55% A at 12 min, 0% A at 13 min, 0% A at 14 min and 20% A at 15 min. A standard calibration was performed using MLG43 (O-BGTRIB; Megazyme) and cellotriose (O-CTR; Megazyme) and quantification was based on the 527 m/z ion [M+Na+]. The chromatogram and spectra were processed using MassLynx software (Waters, Milford, MA, USA).

### Gene expression analysis

Fifteen-day-old wheat plants from the cultivar Titlis were treated, as described above, with 0.5 mM MLG43, 0.5 mM cellotriose or the mock control (0.1% v/v UEP-100 (Croda) and 0.01% v/v Tween 20) for 3 hours. Additionally, for the RNAseq experiment, wheat plants were spray-inoculated with the mock solution, with the wildtype strain and the overexpression mutant line *ZtGH45_OE*, as described above. Samples were collected as in the experiments for quantifying β-glucan oligosaccharide production *in planta* (see above). Three independent biological replicates, each consisting of two second leaves, processed as described in ‘Characterization of *in planta* β-glucan oligosaccharide production’ per treatment were used. Total RNA was extracted using the TRIzol^®^ reagent (Invitrogen, Cat. No. 15596018), treated with DNase I (Qiagen, Cat. No. 792256) and purified using the RNeasy Plant Mini Kit (Qiagen, Cat. No. 74904) according to the manufacturer’s protocol. cDNA was synthesised using the Transcriptor First Strand cDNA Synthesis Kit (Roche Applied Science). Quantitative reverse transcription–polymerase chain reaction (qRT-PCR) experiments were performed as previously described (Meile *et al*., 2020) using the specific primers listed in Supporting Information Table S2. RNA quality check, library preparation (non-stranded polyA enrichment), and subsequent sequencing were performed by Novogene (Beijing, China) using the Illumina HighSeq PE150 platform. About 30 million paired reads of 150 base pairs (bp) per sample were obtained. More than 95% and 1% of the reads were aligned to the *T. aestivum* (GCA_018294505) and the *Z. tritici* ST99CH_3D7 (GCA_900091695) reference genomes (available in Ensembl Plants and Ensembl Fungi), respectively, using the STAR aligner with predetermined parameters (Supporting Information Table S3) (Dobin *et al*., 2013). Uniquely mapped reads were counted using FeatureCounts (Liao *et al*., 2014) in the case of wheat genes and Kallisto (Bray *et al*., 2016) in the case of *Z. tritici* genes with default parameters. Differentially expressed genes (DEGs; Log2-fold change = ± 0.58 with adjusted P value ≤ 0.05) were identified in each comparison using DEseq2 (Love *et al*., 2014). Gene ontology (GO) enrichment analysis was performed using The Gene Ontology Resource tool (https://geneontology.org/) against the biological process annotation dataset. Significantly enriched GO terms were identified with PANTHER Overrepresentation Test using a Fisher’s Exact test with False Discovery Rate correction (p < 0.05).

### Reactive oxygen species (ROS) quantification

Discs (12.6 mm^2^) from second leaves of two-week-old wheat plants of Titlis, Fielder and Paragon cultivars were incubated for 16 h in the dark and at 15 ℃ in 100 µL of a solution containing 150 nM Luminol L-012 (FUJIFILM Wako Pure Chemical Corporation, 120-04891) and 15 µg/mL peroxidase from horseradish (P6782, Sigma-Aldrich). Immediately after, 50 µL of 300 µM MLG43, 300 µM cellotriose, 300 µM hexaacetyl- chitohexaose, 3 µM flg22 or H2O were added, and the luminescence was measured using a Varioskan Lux luminescence reader (Thermo Scientific). ROS production was estimated as relative luminescence units (RLU) over time. At least 8 leaf discs per treatment were used in each experiment. The experiment was performed 3 times independently.

### Stomatal aperture measurement

Wheat epidermal peels isolated from two-week-old plants were incubated in a buffer containing 50 mM KCl, 10 mM MES, 10 mM CaCl2, and pH 6.25 under light for 14 h to open stomata. Subsequently, the epidermal peels were treated with 0.5 mM of MLG43, 0.5 mM cellotriose, 10 µM abscisic acid (ABA) or deionized water for 6 hours. ABA was used as a positive control for inducing stomatal closure (Hsu *et al*., 2021). Images of each stoma were captured with a MikrOkular Full HD camera (Bresser, Germany) attached to a Labophot-2 microscope (Nikon corporation, Japan) using CamLab Lite software. Stomatal width and length were measured using Image J (Schneider *et al*., 2012). The experiment was repeated three times independently.

### Synteny plot and population analyses

To assess the genomic environment of *Zt*GH45, we compared synteny in the 50-kb region surrounding *Zt*GH45 in three reference genomes of *Z. tritici* strains, 3D7, 3D1 and 1E4 assemblies (Plissonneau *et al*., 2018). For this we used a previous annotation of orthologous genes using PoFF, which integrates data on conserved synteny to detect orthologous relationships (Lechner *et al*., 2011). We integrated the transposable element annotation generated by Lorrain et al. (2021) (Lorrain *et al*., 2021) into a synteny plot designed with the R package ggplot2 v3.5.0 (Wickham, 2016).

To investigate the genetic variability found within the *Zt*GH45 coding sequence, we examined the polymorphism found in a Swiss *Z. tritici* population of more than 800 strains (Lorrain *et al*., 2024). We used the variant calling results from Lorrain et al. (2024) and extracted the genomic region corresponding to the ZtGH45 gene (ZtIPO323_110030) (Lapalu *et al*., 2023). We identified the single nucleotide polymorphism (SNP) variants with a predicted non-synonymous effect, using SnpEff (Cingolani *et al*., 2012) with a custom database based on the latest gene annotation of the reference genome IPO323 (Lapalu *et al*., 2023).

### Statistical analysis

Data sets were statistically analysed with Prism 9 software (GraphPad Software, San Diego, California). First, outliers were detected using the ROUT method (Q=1%) and the Gaussian distribution of the data was tested using the Shapiro-Wilk and Kolmogorov-Smirnov tests. Comparisons between two groups were performed by two-tailed *t*-tests. Comparisons between multiple groups normally distributed were performed by parametric ordinary one-way ANOVA followed by Dunnett test or ordinary two-way ANOVA followed by Šídák test. Comparisons between multiple groups non-normally distributed were performed by non- parametric Kruskal-Wallis test together with Dunn’s test.

## Results

### Mixed-linked glucan oligosaccharides are released during *Z. tritici* infection of wheat

To mechanistically explore how *Z. tritici* interacts with the plant cell wall, we monitored the accumulation of β-glucan oligosaccharides (MLG- and cellulose-derived oligosaccharides) during wheat infection. We collected *Z. tritici* (strain 3D7)-infected and non-infected (Mock) wheat leaves at different time points (6, 8 and 10 dpi) and evaluated the presence of β-glucan oligosaccharides with low degree of polymerization (DP; (Supporting Information Fig. S2). We quantified the well-described PTI activators β-glucan trisaccharides (MLG43 and cellotriose) using hydrophilic interaction liquid chromatography in combination with electrospray ionisation-mass spectrometry (HILIC-ESI-MS; Fig. 1a-c and Supporting Information Fig. S2). *Z. tritici* infections led to the release of the MLG-derived oligosaccharide MLG43, which displayed maximum levels upon the appearance of the necrotrophic lesions (10 dpi; Fig. 1b). Additional putative β-glucan oligosaccharides of higher DP were detected (Supporting Information Fig. S2). Remarkably, the levels of the main expected potential products derived from cellulose degradation, cello-oligosaccharides, did not increase during infection (Fig. 1c). To determine the origin of the observed MLG43, we evaluated the presence of β-1,3/1,4-glucan in wheat and *Z. tritici* cell walls. In leaves of 17-day-old wheat plants, β-1,3/1,4- glucan accounted for 0.8% (7.8 ± 0.6 mg/g) of the cell wall (Alcohol Insoluble Residue; AIR), while β-1,3/1,4- glucan was not detected in *Z. tritici* AIR, pointing to wheat as the sole source of the observed release β- 1,3/1,4-glucan oligosaccharides. The results indicate that β-1,3/1,4-glucan-derived oligosaccharides are released from wheat cell walls during *Z. tritici* infection.

**Fig. 1.**
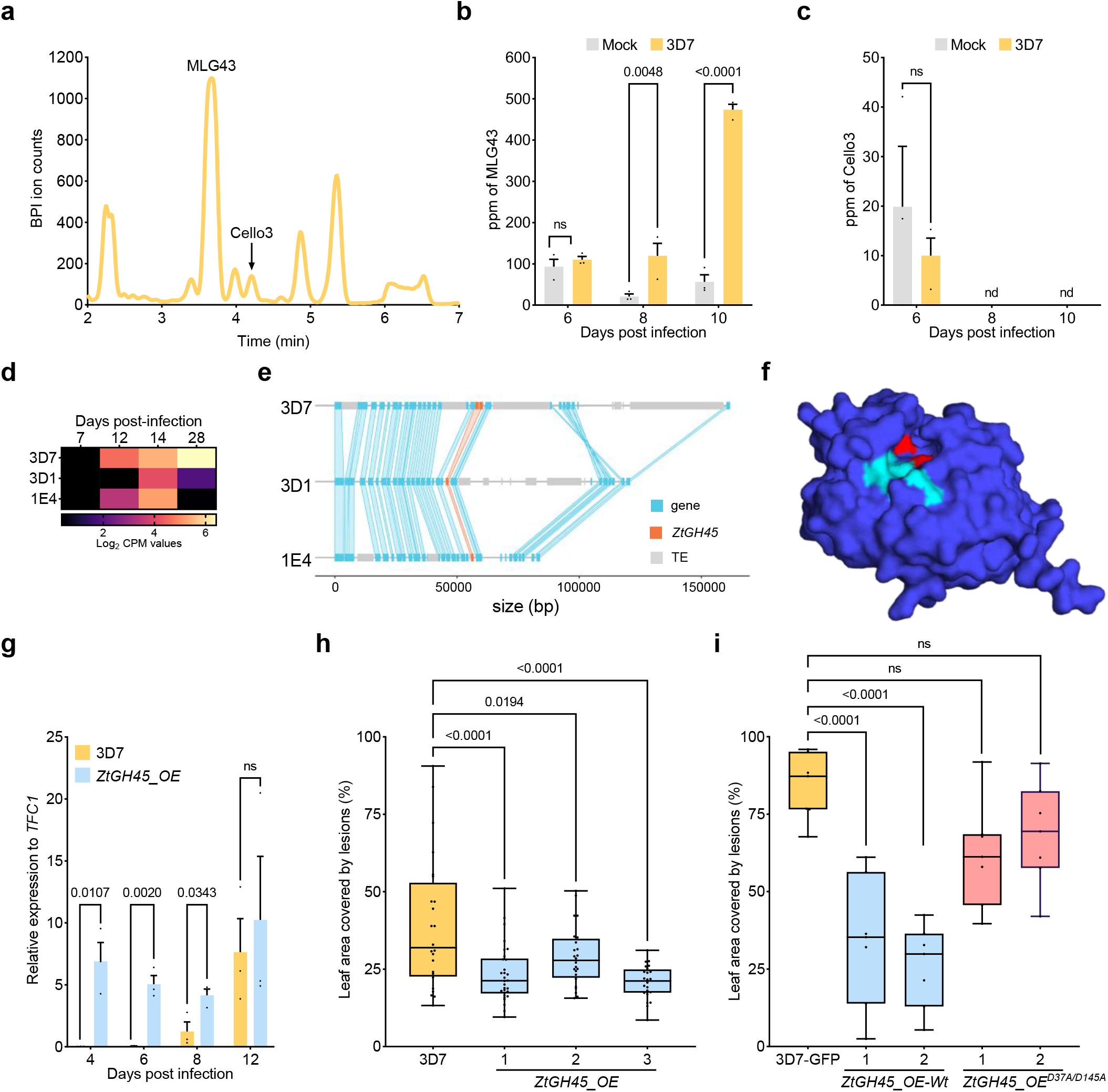
Mixed-linked glucan oligosaccharides are released during *Zymoseptoria tritici* infection and early expression of *ZtGH45* hampers virulence. **a,** Base peak intensity (BPI) chromatogram obtained by hydrophilic interaction liquid chromatography coupled to electrospray ionisation-mass spectrometry (HILIC-ESI-MS) of wheat leaf extracts at 10 days post infection (dpi) with *Z. tritici*. β-ᴅ-cellobiosyl-1,3-β-ᴅ-glucose (MLG43) mean peak and cellotriose (Cello3) peak are labelled. **b**-**c**, Quantification by HILIC-ESI-MS of the mixed linked glucan-derived oligosaccharides MLG43 (**b**) and Cello3 (**c**) released at 6, 8 and 10 days post infection (dpi) with the wild-type *Z. tritici* strain ST99CH_3D7 (3D7). *P* values according to two-way ANOVA followed by Šídák test between plants infected with the 3D7 strain and mock-treated plants are displayed in the plots. Values below detection limit are expressed as not detected (nd). **d**, Expression levels of *ZtGH45* in the *Z. tritici* strains 3D1, 3D7, and 1E4 at 7, 12, 14 and 28 dpi. Data were obtained from a previously published RNA-seq study (NCBI accessions: SRA SRP077418). **e**, Synteny plot comparing region flanking *ZtGH45* between *Z. tritici* strains 3D7, 3D1 and 1E4. Genes are represented by blue blocks, *ZtGH45* is shown in red blocks and transposable elements (TEs) are represented by grey blocks. Collinear sequences between the two strains are shown in blue and red. **f**, Surface representation of *Zt*GH45 structure model obtained using Alphafold2 (pLDDT: 92.7; ColabFold v1.5.5). Residues described to mediate oligosaccharide-binding (cyan), and residues in the catalytic site (red), according to the *Nc*GH45 homologue protein from *N. crassa,* are coloured. **g**, Expression pattern of *ZtGH45* in the wild-type strain 3D7 (yellow bars) and in the lines constitutively overexpressing *in locus ZtGH45* (ZtGH45-overexpression line in locus, *ZtGH45_OE*, blue bars) during wheat (cultivar Titlis) infection at 4, 6, 8, and 12 dpi. Values shown are relative to the *Z. tritici* housekeeping gene *TFC1*. *P* values according to a two- tailed *t*-test between plants infected with the *ZtGH45_OE* line and the 3D7 strain are displayed in the plots. **h**, Percentage of leaf area covered by lesions produced by 3D7 (control, yellow bar) and *ZtGH45_OE* (blue bars) strains at 13 dpi on wheat plants of cultivar Titlis. The experiment was performed three times with line #3 and we obtained similar results. **i**, Percentage of leaf area covered by lesions produced in Titlis wheat plants by 3D7-GFP (control, yellow bar) and lines ectopically overexpressing the *ZtGH45* wild-type version (*ZtGH45_OE-WT*; blue bars) or *Zt*GH45 with the catalytic site mutated (*ZtGH45_OE*^D37A/D145A^; red bars) at 13 dpi. In both **h** and **i**, the *P* values according to a one-way ANOVA followed by Dunnett test between plants infected with the mutant lines and the control strain are displayed in the plots.

We next aimed to identify the CAZyme(s) secreted by *Z. tritici* which led to the production of MLG43. We hypothesised that enzymes leading to MLG43 accumulation should have a predicted glucanase activity (EC 3.2.1.4) and be highly expressed at the beginning of the necrotrophic phase. Of the 27 *Z. tritici* genes encoding GHs with a predicted glucanase activity (from the families GH5, GH7, GH10, GH12, GH45 and GH51), one (*ZtGH45*/*Zt3D7_G10118*) displayed very low expression levels at 7 dpi and when grown under axenic conditions, was highly induced at the onset of the necrotrophic phase (12 dpi), and exhibited expression levels that remained high during the necrotrophic phase (Supporting Information Fig. S3;Palma-Guerrero et al., 2017; Francisco et al., 2019). Furthermore, we showed that *ZtGH45* displayed a similar expression pattern in other *Z. tritici* strains (3D1 and 1E4), reaching maximum levels at the switch to the necrotrophic phase (Fig. 1d). Previous work demonstrated that *ZtGH45* resides in a transposable element-rich region and its expression is regulated by chromatin changes involving H3K27me3 and H3K9me3 in the strain 3D7 (Meile *et al*., 2020). We showed that *ZtGH45* was in the proximity of transposable elements in the three investigated strains (3D7, 3D1 and 1E4), which suggests a potential epigenetic regulation of *ZtGH45* in other *Z. tritici* strains (Fig. 1e). ZtGH45 harbours 6 cysteine bridges, as shown in an Alphafold2 model, is highly polymorphic, and was previously shown to be under significant diversifying selection (Fig.1f; Supporting Information Fig. S4b; Brunner et al., 2013). Despite the high number of polymorphic residues of ZtGH45, the critical residues for GH45 activity (D37 and D145), as reported in *Neurospora crassa* and *Thermothielavioides terrestris* (Gao *et al*., 2017; Kadowaki & Polikarpov, 2019), were shown to be conserved in all the investigated strains of *Z. tritici*, suggesting a potential β-1,4-glucanase activity of ZtGH45 (Supporting Information Fig. S4a).

### Impact of constitutive *ZtGH45* expression on *Z. tritici* virulence

We functionally characterised *Zt*GH45 by obtaining overexpression lines in which we substituted the native promoter by the constitutive *gdpA* promoter (*ZtGH45*-overexpression line *in locus*, *ZtGH45_OE*). In the wild- type control, *ZtGH45* is not expressed under axenic conditions and has shallow expression levels at the early stages of the infection, as previously reported (Meile *et al*., 2020; Suarez-Fernandez *et al*., 2023) (Fig. 1g and Supporting Information Fig. S5a). In contrast, in the *ZtGH45_OE* line, *ZtGH45* expression levels remained high throughout the *Z. tritici* infection cycle (Fig. 1g). The three tested *ZtGH45_OE* lines showed reduced virulence in the cultivar Titlis in comparison to the wild-type strain, as reflected by a reduction in the percentage of leaf area covered by lesions, and by impaired reproduction, as shown by diminished pycnidia density (Fig. 1h-i and Supporting Information Fig. S5b). This effect was not cultivar-specific since we observed a similar effect in the cultivar Drifter (Supporting Information Fig. S5c-d). *ZtGH45* expression levels were not significantly different between the wild-type and *ZtGH45_OE* lines during the necrotrophic phase (Fig. 1g). These data suggested that the reduced virulence phenotype observed in the *ZtGH45_OE* lines was attributable solely to the misexpression of the gene at the early stages of infection. To rule out the possibility that the virulence phenotype resulted from altered growth of the mutant, we conducted an *in vitro* growth analysis in minimal medium supplemented with different carbon sources (sucrose, fructose, carboxymethyl cellulose (CMC) and barley β-glucan) and in yeast malt sucrose (YMS) medium in the presence of different abiotic stresses (sorbitol, NaCl, H2O2, Calcofluor white, and 28 ℃). No differences in growth were observed between the *ZtGH45_OE* and the wild-type lines (Supporting Information Fig. S6). These results demonstrate that misexpression of *ZtGH45* hampered the ability of *Z. tritici* to colonise the host.

We next explored whether the reduced virulence of *ZtGH45_OE* was due to the recognition of the *Zt*GH45 protein itself or due to the recognition of the released oligosaccharides from the wheat cell wall by the activity of *Zt*GH45. With this aim, we obtained a mutant line that overexpressed an enzymatically inactive version of *Zt*GH45 (gdpA::*Zt*GH45^D37A/D145A^; *Zt*GH45^D37A/D145A^_OE). Remarkably, we observed that ectopic overexpression of the catalytically dead version of *Zt*GH45 (*Zt*GH45^D37A/D145A^_OE) did not impair virulence, while ectopic overexpression of the wild-type version of *Zt*GH45 hindered *Z. tritici* progression (Fig. 1i and Supporting Information Fig. S7). The results demonstrate that the phenotype of *ZtGH45_OE* is exclusively attributable to the catalytic activity of *Zt*GH45, and that the protein itself is not recognized by the wheat immune system.

### Constitutive expression of *ZtGH45* leads to enhanced release of MLG-derived oligosaccharides

We further explored whether *ZtGH45_OE* lines exhibit an enhanced capacity to hydrolyse β-glucans. We cultivated *Z. tritici* wild-type and *ZtGH45_OE* lines in the presence of 0.5% (w/v) β-1,3/1,4-glucan or β-1,4-glucan (CMC). β-glucan oligosaccharides quantification of the culture filtrates after 96 h of growth revealed that the *ZtGH45_OE* line significantly released more β-glucan oligosaccharides than the wild-type strain when employing β-1,3/β-1,4-glucan as a substrate (39%, 51%, and 66% for each independent *ZtGH45_OE* line and 15% for the wild-type; % w/w of the initial material) and these levels were similar to those generated by incubating MLG with the enzyme cellulase (Fig. 2a and Supporting Information Fig. S8a). When using CMC as a substrate, we detected the release of β-1,4-glucan oligosaccharides (0.2% of the initial material; w/w) only in the *ZtGH45_OE* lines, and not in the wild-type strain. (Fig. 2b and Supporting Information Fig. S8b). To bring the experimental conditions closer to the *in vivo* wheat environment, the wild-type and a mutant line were grown in the presence of wheat cell wall fraction (AIR) obtained from 17-day-old seedlings. β-1,3/1,4-glucan and β-1,4-glucan-derived oligosaccharides were released by both the wild-type and the *ZtGH45_OE* line cultivated in this wheat AIR wall fraction, but the *ZtGH45_OE* line released 1.8-fold higher levels of β-glucan oligosaccharides than the wild-type strain (Fig. 2c). Oligosaccharides released by *ZtGH45_OE* from β-1,3/1,4- glucans were further characterised using high-performance anion-exchange chromatography with pulsed amperometric detection (HPAEC-PAD). *ZtGH45_OE* exhibited an enhanced production of MLG43 and other β-glucan oligosaccharides of different degree of polymerization compared to the wild-type strain (Fig. 2d and Supporting Information Fig. S8c). These findings corroborate the β-1,4-glucanase activity of *Z. tritici Zt*GH45, highlighting its ability to hydrolyse β-1,3/1,4-glucan from the wheat cell wall, leading to the release of β- glucan oligosaccharides.

**Fig. 2.**
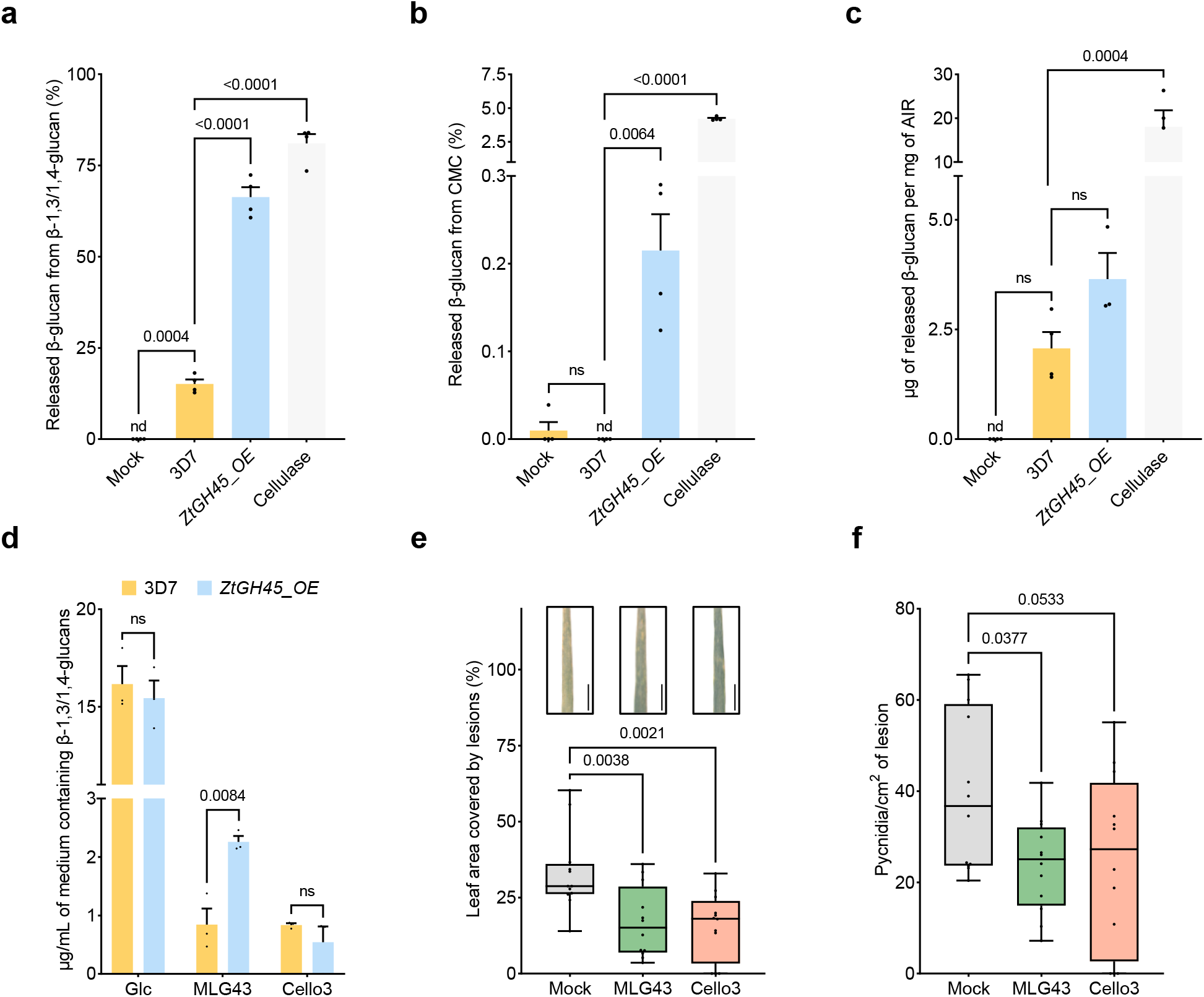
*ZtGH45* overexpression leads to an enhance degradation of wheat cell wall β-glucan polysaccharides and to the release of oligosaccharides that trigger resistance against *Zymoseptoria tritici*. **a-c**, β-glucan oligosaccharides released by *Z. tritici* wild-type (3D7) and *ZtGH45-*overexpression (*ZtGH45_OE*) lines grown in Vogel’s minimal medium supplemented with 0.5% (w/v) fructose and with either 0.5% (w/v) barley mixed-linked glucan polysaccharide (β-1,3/1,4-glucan) (**a**), 0.5% (w/v) carboxymethylcellulose sodium salt (CMC) (**b**) or 0.5% (w/v) wheat alcohol insoluble residue (AIR) (**c**). Pure endo-1,4-β-D-glucanase (cellulase; 2.5 U/mL) and water (mock) treatment were used as positive and negative controls, respectively. Oligosaccharides were quantified after 96 hours (**a**-**b**) or 120 hours of growth (**c**). Bars shown in **a**-**c** panels represent the mean of four biological replicates and the error bars represent the standard error of the mean. *P* values according to one-way ANOVA followed by Dunnett test between mock, *ZtGH45_OE* line or cellulase treatment and 3D7 strain are displayed in the plots. The experiments were performed two times, and we obtained similar results. Values below detection limit are expressed as not detected (nd). **d**, β-glucan trisaccharides (β-ᴅ-cellobiosyl-1,3-β-ᴅ-glucose; MLG43 and cellotriose; Cello3) and glucose (Glc) released by *Z. tritici* wild-type (3D7) and *ZtGH45-*overexpression (*ZtGH45_OE*) lines after 96 hours of growth in β-glucan- supplemented media. The oligosaccharides were quantified by high-performance anion-exchange chromatography with pulsed amperometric detection (HPAEC-PAD). *P* values according to a two-tailed *t*-test between plants infected with the *ZtGH45_OE* and 3D7 strain are displayed in the plot. Bars represent the mean of three biological replicates and the error bars represent the standard error of the mean. **e**-**f**, *Z. tritici* strain 3D7 virulence measured as percentage of leaf area covered by lesions (**e**) and pycnidia density (pycnidia/cm^2^ of lesion) (**f**), at 14 days post infection (dpi) in Titlis wheat plants pre-treated with 1 mL of 0.5 mM β-ᴅ-cellobiosyl-1,3-β-ᴅ-glucose (MLG43; green bar) or 0.5 mM cellotriose (Cello3; red bar) 24 hours before infection. *P* values according to one-way ANOVA followed by Dunnett test between the oligosaccharide treatments and the mock control are displayed in the plots. One representative picture of each treatment is shown in **e**. Scale bars, 1 cm. An additional replicate of the results presented in **e** is shown in Supporting Information Fig. S7 d.

### Mixed-linked glucan oligosaccharides trigger wheat immunity

We hypothesised that the impaired virulence of the *ZtGH45_OE* lines was due to an early *Zt*GH45-dependent release of β-glucan oligosaccharides. To test this hypothesis, we determined whether these *Zt*GH45-derived oligosaccharides act as plant immunity-activating molecular patterns. MLG43 and cellotriose treatment of wheat leaves 24 hours before inoculation with the wild-type strain resulted in a reduction in symptoms and pycnidia formation compared to mock-treated plants in the cultivar Titlis after 16 days of infection (Fig. 2e-f; Supporting Information Fig. S8d). These results suggest that these β-glucan oligosaccharides prime wheat plants against pathogen attack. To determine whether these cell wall-derived oligosaccharides induced transcriptional reprogramming in wheat, including the activation of defence-related genes, we performed a comparative transcriptomic analysis. We analysed the wheat transcriptome three hours after treatment of plants with MLG43, cellotriose or the mock solution. In the principal component analyses, MLG43-induced transcriptional changes compared to mock were primarily captured by PC1, explaining 88% of the total variance, while cellotriose-associated changes were delineated by PC2, which accounted for barely 4% of the total variance (Fig. 3a). This observation indicates that MLG43 triggers a more pronounced transcriptional reprogramming in wheat plants compared to cellotriose. Accordingly, when comparing the transcriptomic profiles of treated and control samples, we identified a total of 26 and 3997 differentially expressed genes (DEG; fold change (Log2) ± 0.58, P adjusted value ≤ 0.05), upon treatment with cellotriose and MLG43, respectively, indicating that these oligosaccharides trigger different transcriptomic responses (Supporting Information Tables S4-S5). In accordance with its capacity to act as a DAMP, the 3173 up-regulated genes upon MLG43 treatment were enriched in GO terms related to disease resistance responses (“innate immune response” (GO:0045087), “defence response to other organism” (GO:0098542), and “transmembrane receptor protein serine/threonine kinase signalling pathway” (GO:0007178) (Supporting Information Table S6) and included 178 genes encoding for defence-related proteins, consisting of homologues of 22 chitinases, and 149 receptor-like kinases, including CERK1, CEBiP (chitin elicitor-binding protein), LYK5 (LysM motif receptor kinase 5), two LRR-MAL receptor kinases, 21 LRR receptor kinases, six wall-associated receptor kinases (WAK), and 23 LecRKs (Supporting Information Table S7, highlighted in green). Remarkably, cell wall- related GOs (GO:0071554 and GO:0044036) were also significantly enriched in the genes induced by MLG43 (Supporting Information Table S6). Overall, the results suggest that MLG43 treatment leads to a transcriptional reprogramming involving the induction of an immune response and cell wall remodelling and suggest an enhanced response in wheat to MLG43 compared to cellotriose.

**Fig. 3.**
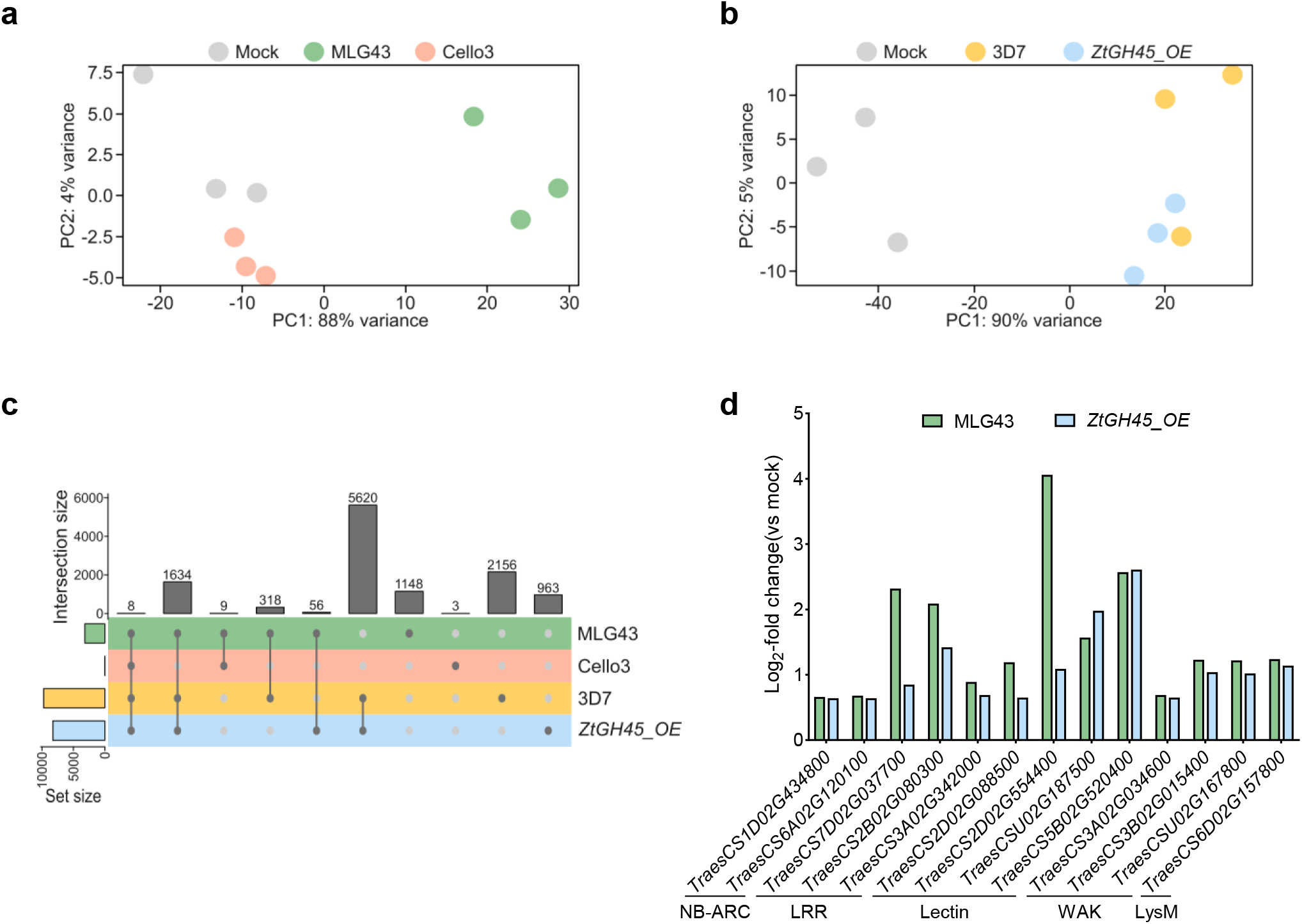
Mixed-linked glucan oligosaccharides trigger a defence response. **A**, Principal component analysis (PCA) of wheat transcriptomes 3 hours after treatment with 0.5 mM MLG43 (green dots), 0.5 mM Cello3, (red dots) or mock conditions (blue dots). **B**, PCA of wheat transcriptome 24 hours after spray inoculation with the wildtype strain 3D7, with the mutant line *ZtGH45_OE* or with the mock solution. **C**, UpSet plot showing the overlapped of upregulated genes between comparisons. Horizontal bars represent the number of DEGs for each comparison. The intersection size of up-regulated genes is shown in the vertical bar plot. **C**, Log2 fold change levels of potential resistance genes up-regulated by MLG43 and the infection with *ZtGH45_OE*. The potential resistance genes containing a Lectin domain, a LysM domain and an LRR domain are show. Additionally, the two wall-associated kinases (WAK) and the two NB-ARC-containing proteins are represented.

In parallel, we also investigated the transcriptional response of wheat plants to infection with the line constitutively expressing *ZtGH45* (*ZtGH45_OE*). Wildtype (3D7), *ZtGH45_OE*-infected, and non-infected control plants were collected at an early time point of the infection (6 dpi) when macroscopic symptoms were not observed (Bernasconi *et al*., 2023; Alassimone *et al*., 2024) and the wild-type strain did not express *ZtGH45*. The principal component analysis highlighted that infection with the wildtype and the *ZtGH45_OE* lines led to changes in the transcriptome compared to the mock, explaining 90% of the total variance (Fig. 3b). In total, 20451 and 17375 wheat genes were differentially expressed upon infection with the wild-type and the overexpression line, respectively, compared to the control plants, of which 14940 genes were shared and 2435 were specific to *ZtGH45_OE* infection (Supporting Information Tables S4-S5). Of the total up- regulated genes upon infection, we identified 1019 genes (12.3%) that were induced only upon *ZtGH45_OE* infection (Fig. 3c and Supporting Information Table S8). These genes specifically induced upon *ZtGH45_OE* infection were enriched in “regulation of β-glucan biosynthetic process” (GO:0032951), “response to other organisms” (GO:0051707) and “phosphorylation” (GO:0016310) GO terms (Supporting Information Table S9). Among these genes, two encoded for 1,3-β-glucan synthases, 25 for receptor-like kinases, and one for a receptor kinase with a lectin-domain (Supporting Information Table S8, highlighted in green). From the fungal side, only seven genes were differentially regulated in the *ZtGH45_OE* line compared to the wild-type strain (Supporting Information Tables S10-S11). We further evaluated if the transcriptomic profiles of the *Z. tritici* and MLG43 treatments overlapped. We found that more than 60% (2016) of the genes up-regulated by MLG43 were also induced upon *Z. tritici* infection (Fig. 3c). Of these, 56 genes were specifically regulated by *ZtGH45_OE* and not by the 3D7 wild-type strain, including 13 genes involved in stress perception and signalling (Table 1; Fig. 3d). Overall, we identified a set of defence-related genes induced after treatment of wheat with MLG43 and infection with *ZtGH45_OE*.

**Table 1.**
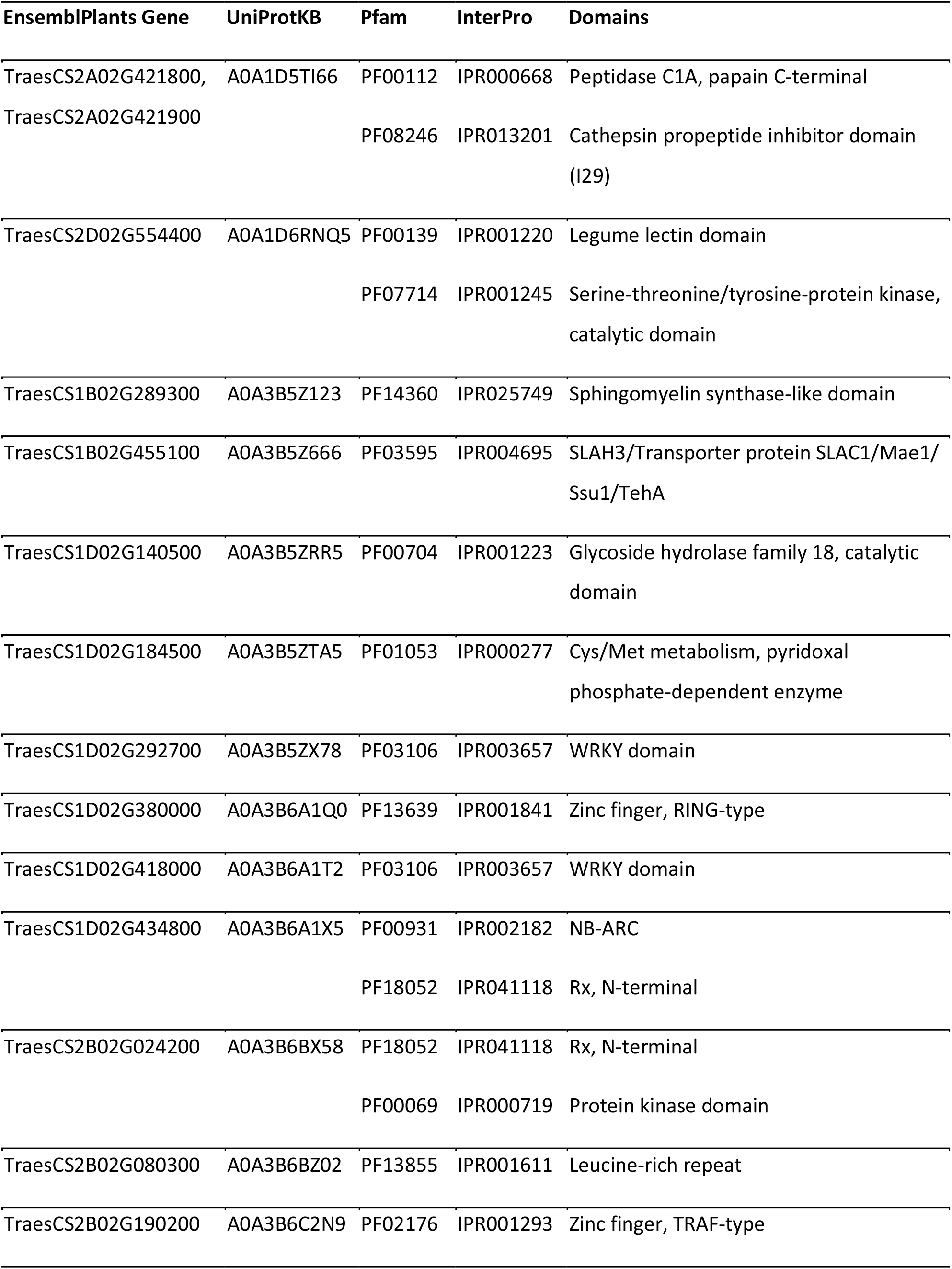
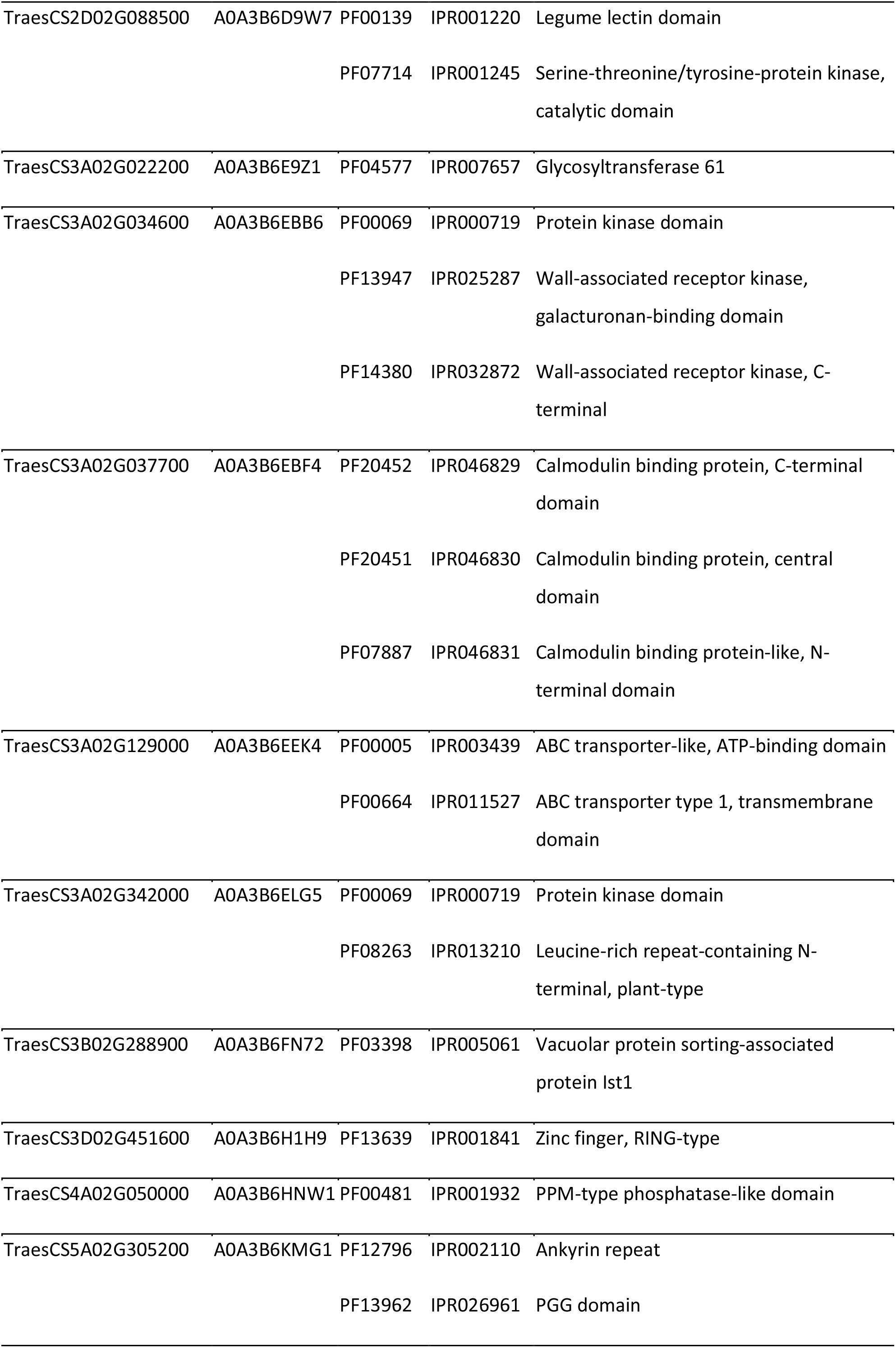
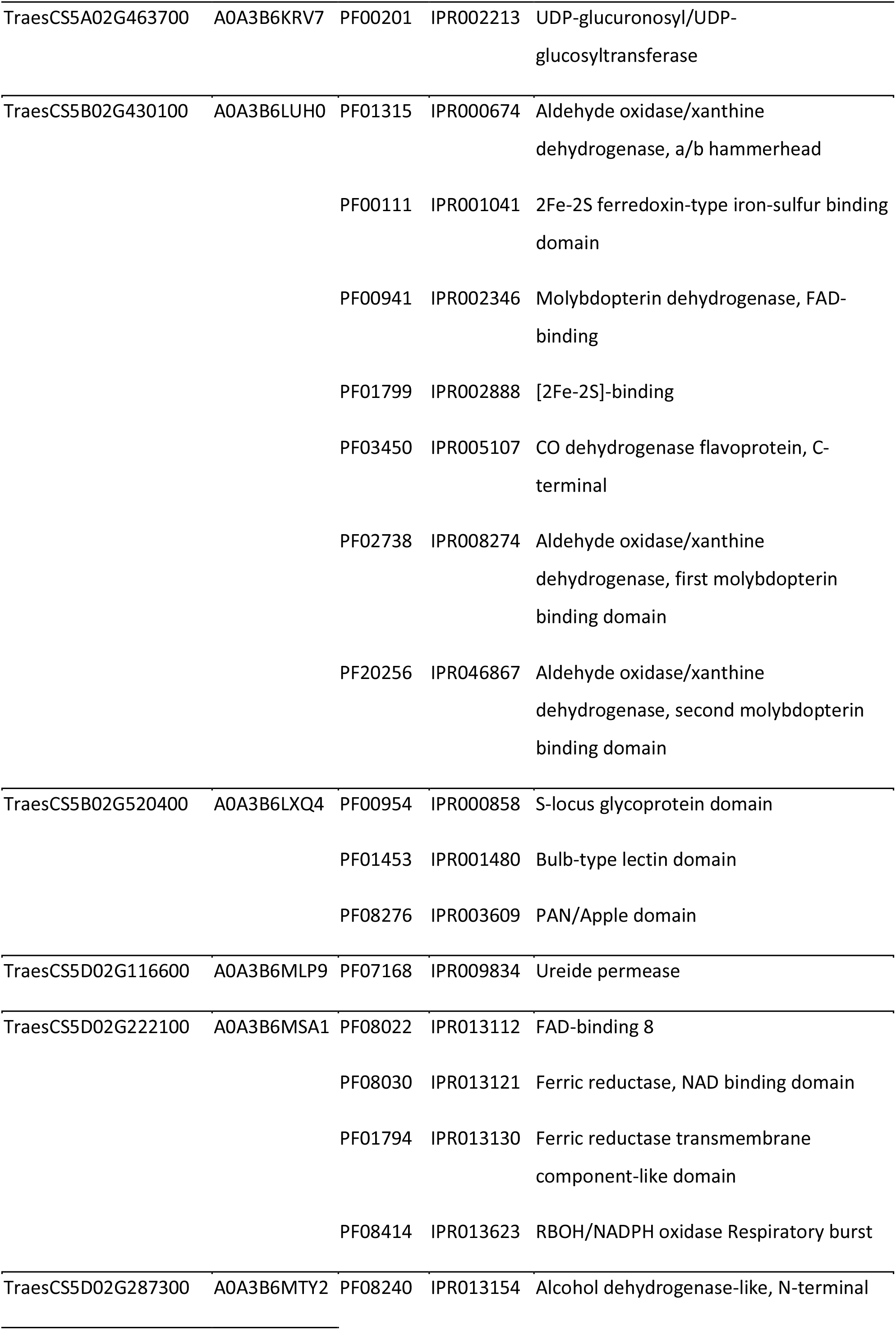
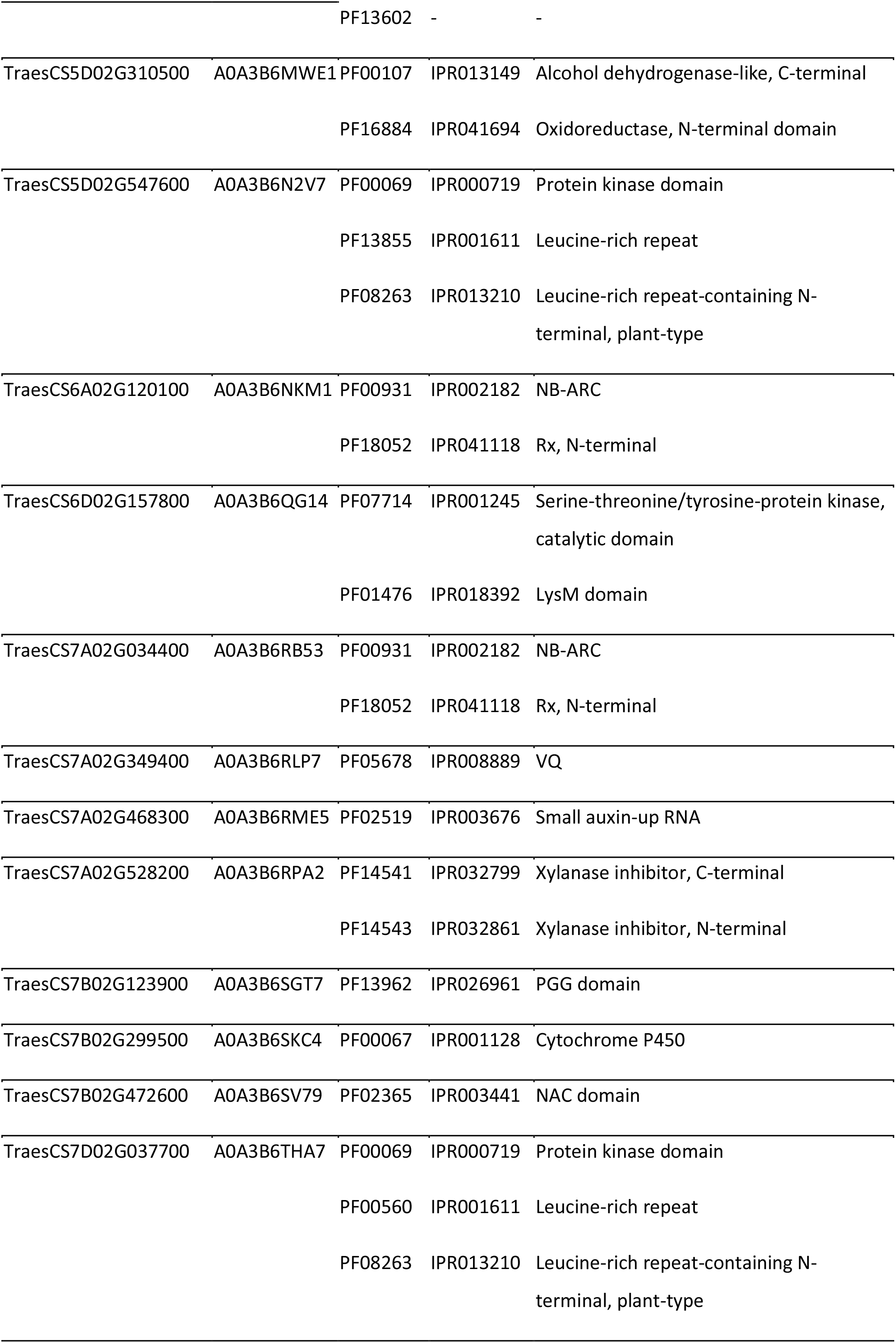
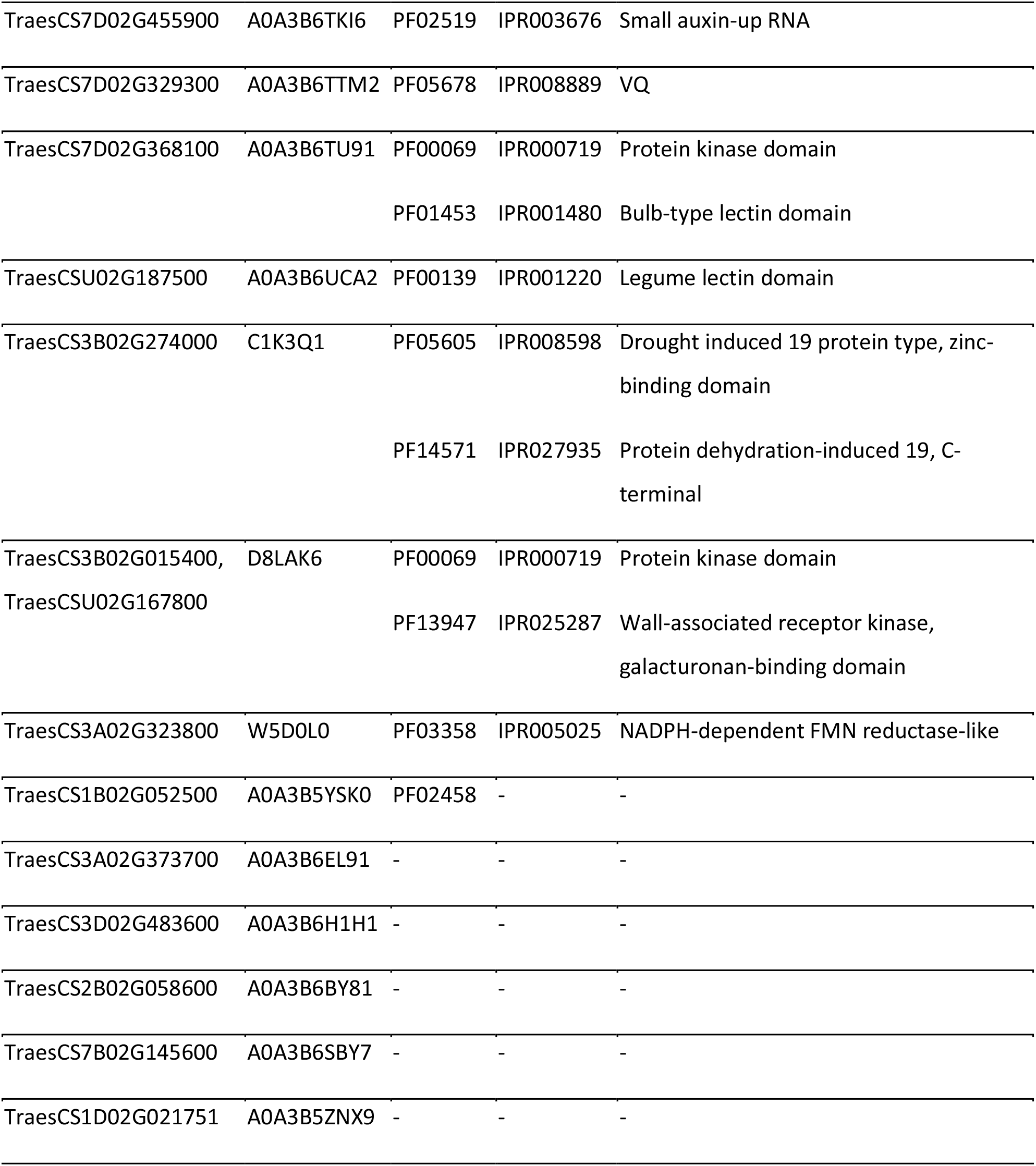
Genes up-regulated in wheat upon infection with *Zymoseptoria tritici Zt*GH45-overexpression line (*Zt*GH45_OE; 6 days post infection), but not with the wildtype strain, and upon treatment with 1 mL of 0.5 mM of MLG43 (3 h after treatment).

### MLG43 triggers a ROS burst and stomatal closure

Since five wheat respiratory burst oxidase (*RBOH*) genes were upregulated upon MLG43 treatment (Fig. 4a), we investigated ROS production upon treatment with this elicitor in cultivar Titlis. We observed that wheat leaves underwent a ROS burst upon treatment with MLG43 (Fig. 4b-e). Remarkably, ROS burst was also observed in cultivars Fielder and Paragon, indicating that MLG43 triggers this immune response regardless of the wheat cultivar used (Fig. 4d-e). In contrast, wheat treated with cellotriose did not undergo ROS accumulation (Fig. 4b-c). Additionally, the transcriptomic analysis revealed an upregulation of 15 genes involved in stomatal closure regulation, in addition to the five *RBOHs* (Kwak *et al*., 2003) indicated above. These 15 genes included genes encoding for homologues of two SLAH3 (Liu *et al*., 2019), eight plant glutamate receptor (Kong *et al*., 2016), four calcium-dependent protein kinases (CPKs; Geiger et al., 2010; Scherzer et al., 2012; Brandt et al., 2012), and one CBL-interacting protein kinase (CIPK; Fig. 3b; Förster et al., 2019), suggesting that MLG43 might regulate stomatal closure. Since stomata have been shown to play a pivotal role in plant resistance and are the major entry gate for *Z. tritici* to colonise the wheat apoplast (Battache *et al*., 2022; Alassimone *et al*., 2024), we investigated whether *Zt*GH45-released cell wall oligomers regulate stomatal movements by treating wheat leaves with MLG43 and cellotriose and assessing stomatal opening. Remarkably, we observed that 6 hours after MLG43 treatment, wheat plants closed the stomata, similar to the response of plants to treatment with abscisic acid (ABA; Fig. 4f; Hsu et al., 2021), used as positive control). In contrast, stomata remained open upon cellotriose and water treatment (Fig. 4f). These results indicate that the recognition of MLG43 leads to ROS accumulation and subsequent stomatal closure in wheat plants.

**Fig. 4.**
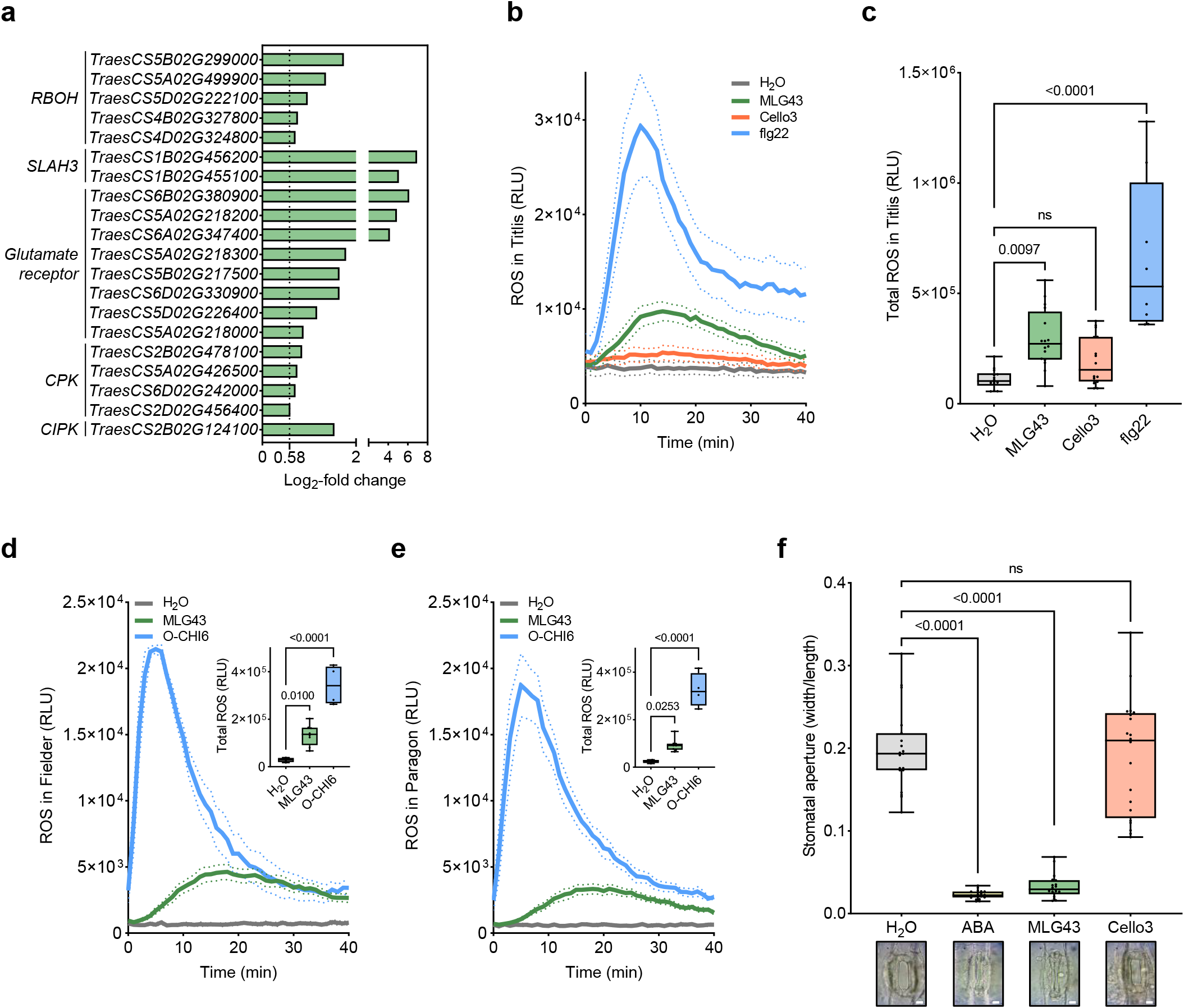
Mixed-linked glucan oligosaccharides trigger ROS burst and stomatal closure. **a**, Expression levels of stomatal immunity-related genes in wheat plants treated with 1 mL of MLG43. Log2- fold change compared to mock are represented. **b**, Reactive oxygen species (ROS) production in wheat leaf discs of cultivar Titlis upon treatment with 100 µM MLG43 or cellotriose (Cello3). measured as relative luminescence units (RLU) over time. 1 µM flagellin-derived 22-amino-acid epitope (flg22) was used as a positive control, respectively. Dashed lines represent the standard error of the mean. **c**, Total ROS production estimated as cumulative RLUs over 40 min in cultivar Titlis. These experiments were repeated twice and gave similar results. (**d-e**) ROS production, estimated as RLUs, in wheat leaf discs of cultivar Fielder (**d**) and Paragon (**e**) upon treatment with 100 µM MLG43 or 100 µM hexaacetyl-chitohexaose (O-CHI6). Total ROS production is shown in the right upper corner. In b-e, water was used as negative control and *P* values according to one- way ANOVA followed by Dunnett test between oligosaccharide- or flg22-treatment and H2O are displayed in the plots. **f**, Stomatal closure in response to MLG43. Boxplots represent the width/length ratio of the stomatal pore, as a proxy for stomatal aperture, in wheat epidermal peels treated with 0.5 mM of MLG43 or 0.5 mM Cello3 for 6 hours. Water and 10 µM ABA were used as negative and positive controls, respectively. *P* values according to Kruskal-Wallis test followed by Dunn test between MLG43, Cello3 or ABA treatment and H2O are displayed in the plots. This experiment was repeated twice and gave similar results. Representative images of stomata exposed to 10 µM ABA, 0.5 mM MLG43 or 0.5 mM Cello3 for 6 hours are shown. Scale bars, 10 μm.

## Discussion

The plant cell wall plays a central role in mediating plant-pathogen interactions since it acts as a defensive barrier against pathogen invasion and serves as a source of nutrients for pathogens (Molina *et al*., 2024b). Pathogens secrete an arsenal of CAZymes (including CWDEs) during their infection cycle (Brunner *et al*., 2013; Bradley *et al*., 2022) to modify and hydrolyse plant cell wall carbohydrate polymers, presumably to surmount the cell wall as a physical barrier and to acquire nutrients (Bradley *et al*., 2022). As a result of CAZyme activity, cell wall-derived oligosaccharides are released and frequently detected by the host as DAMPs, triggering immune responses (Bacete *et al*., 2018; Molina *et al*., 2024b). In this work, we functionally characterised a glycoside hydrolase from the family 45 from the fungal plant pathogen *Z. tritici*, *Zt*GH45. Early expression of *ZtGH45* leads to an enhanced release of plant cell wall-derived MLG oligosaccharides, which are recognized by the wheat host as immunity-activating molecular patterns. These results suggest that *Z. tritici* tightly regulates *ZtGH45*, which is expressed only during the necrotrophic phase, to delay the release of MLG oligosaccharides and prevent early recognition by the host. Our experiments demonstrate that tight regulation of pathogen CWDEs is necessary to control the release of elicitors that trigger plant resistance.

The plant cell wall is composed of several polysaccharides, with cellulose, a β-1,4-glucose polymer, being the main component (Carpita & McCann, 2000). Cellulose forms paracrystalline structures known as microfibrils that are cross-linked by non-cellulosic polysaccharide that, in the case of Poaceae species manly consist of β- 1,3/1,4-glucan (MLG), and heteroxylans (Burton & Fincher, 2014). *Z. tritici* infection leads to changes in the cell wall composition, featuring an increase of oligosaccharides derived from β-1,3/1,4-glucan, rather than from cellulose. We hypothesised that this might be due to the higher accessibility of MLG in the plant cell wall compared to β-1,4-glucan, which forms the cellulose microfibrils (Zoghlami & Paës, 2019). Additionally, we cannot discard the possibility that cello-oligosaccharides with a low degree of polymerization might be targeted by degrading enzymes, including cellodextrin oxidases, from the plant, as reported in Arabidopsis (Costantini *et al*., 2023), and are therefore not detected. The released MLG-derived trisaccharide is immunogenic and protects wheat plants against *Z. tritici*, similar to what has been described for other plant species, including rice and *A. thaliana* (Rebaque *et al*., 2021; Yang *et al*., 2021). In these two cases, perception of MLG43 is mediated by LysM receptor kinases, and OsLecRK1 and AtLRR-MAL RKs (Martín-Dacal *et al*., 2023; Yang *et al*., 2021; Rebaque *et al*., 2021). The mechanisms mediating MLG43 perception in wheat remain unknown, but MLG43 detection leads to the activation of several defence-related genes, including three LysM-containing receptors, two LRR-MAL RK, 21 LecRKs, and six genes encoding WAKs. The upregulation of these putative receptor-encoding genes in wheat upon MLG43 treatment suggests their potential contribution to the perception of this elicitor. In addition to the induction of defence-related genes, MLG43 triggers the accumulation of ROS and stomatal closure in wheat, which probably limits penetration by *Z. tritici* and potentially accounts for the observed protection capacity of MLG oligosaccharides.

We hypothesise that during colonisation pathogens, including *Z. tritici*, seek to minimise the generation of cell wall-derived oligosaccharides acting as elicitors to enhance the infection process. Accordingly, MLG- oligomer accumulation increases at the later stages of the infection when the fungal hyphae grow quickly, and necrotrophic symptoms first appear. We suggest that MLG43-triggered immune responses at this late stage, when the pathogen grows profusely in the apoplast, cannot limit further development of the infection, while the infection would be effectively hindered if MLG43 was produced at an earlier stage of the infection. Thus, the production of MLG oligosaccharide-releasing enzymes (as *Zt*GH45) is repressed in *Z. tritici* during the early colonising stages of the infection as a mechanism of host evasion. In *M. oryzae*, MoCel12 hydrolyses MLG and releases MLG oligosaccharides and, as we observed for *ZtGH45*, the expression levels of *MoCel12* are regulated to prevent early detection of the pathogen (Yang *et al*., 2021). As described for other effector genes, *ZtGH45* is located in a TE-rich region of the genome which is enriched in heterochromatic marks (H3K27me3 and H3K9me3; Fig. 1e; Meile et al., 2020). We previously demonstrated that *ZtGH45* is epigenetically silenced during the early stages of the infection and only de-repressed during a later phase (Meile *et al*., 2020; Suarez-Fernandez *et al*., 2023). Our results demonstrate that preventing early expression of this effector gene is key for pathogen virulence, highlighting the relevance of chromatin remodelling for pathogen virulence. We postulate that *Z. tritici* pathogenicity balances the release of CWDEs to enable pathogen colonization and nutrient acquisition while preventing their early recognition through tight regulation of CWDEs.

Biotechnological strategies for crop protection based on the capacity of plant cell wall-derived oligosaccharides to increase plant resistance to pathogens have recently gained interest (Molina *et al*., 2024a). Notably, pre-treatment of plants with MLG oligosaccharides (especially MLG43) can protect different crops such as pepper, rice and tomato against bacterial and fungal diseases (Rebaque *et al*., 2021; Yang *et al*., 2021). The ability to obtain these active oligosaccharides from plant-based industrial waste and plant biomass would contribute to circular economy and enable more sustainable agriculture (Rebaque *et al*., 2023). The results shown here illustrate the potential to apply this technology to wheat crops.

In conclusion, β-1,3/1,4-glucan oligosaccharides produced during *Z. tritici*-wheat interactions serve as danger signals triggering wheat immunity that leads to the production of ROS, stomatal closure and a transcriptional response (Fig. 5). We propose that coevolution limited the expression of the β-glucan-degrading enzyme ZtGH45, and probably other CAZymes, to late necrotrophic stages when the pathogen has already colonised the apoplastic space, melanized pycnidia are being formed and the MLG-triggered plant immune response may not effectively interfere with the pathogen progression.

**Fig. 5.**
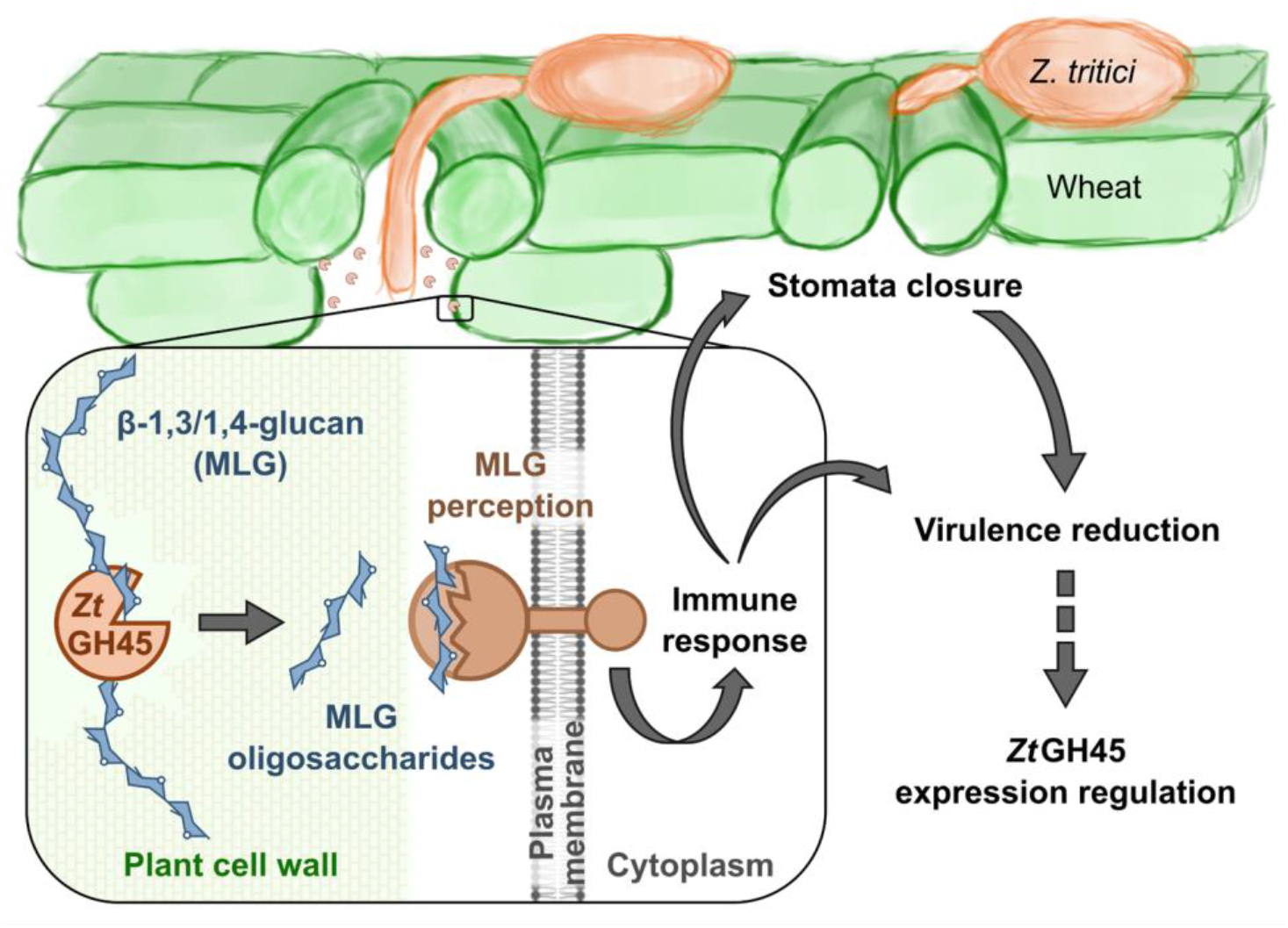
Proposed model for the function of *Zt*GH45 during *Zymoseptoria tritici* infection of wheat plants. Mixed-linkage glucan (β-1,3/1,4-glucan; MLG) oligosaccharides is released during *Z. tritici* infection from wheat cell walls at the onset of the necrotrophic phase. MLG-derived oligosacharides are perceived by wheat and trigger an immune response, featuring an induction of defence-related genes, production of reactive oxygen species and stomatal closure, which hinders the progression of *Z. tritici*. We hypothesise that *Z. tritici* has evolved to express *ZtGH45* at late stages of the infection to prevent the induction of early immune responses by the host.

## Acknowledgments

This work was supported by grant PID2019-108693RA-I00 funded by MCIN/AEI/ 10.13039/501100011033 and RYC2018-025530-I grant from the Spanish Ministry of Science, Innovation, and Universities to ASV and grant PID2021-126006OB-I00 to AM, funded by MCIN/AEI/ 10.13039/501100011033 and by “ERDF A way of making Europe’’ and grant "Severo Ochoa Program for Centres of Excellence in R&D (grant SEV-2016-0672 and CEX2020-000999-S (2022-2025)) funded by MCIN/AEI/ 10.13039/501100011033. CCL and LM have been financially supported as postdoctoral researchers by CEX2020-000999-S (2022-2025) and CNS2023-144037 (funded by MICIU/AEI/10.13039/501100011033 and by EU NextGenerationEU/PRTR), respectively. DR was the recipient of Industrial PhD fellow from Madrid Regional Government (IND2017/BIO-7800) and a “Margarita Salas” postdoctoral fellowship European Union-NextGeneration EU UP2021-035 (RD 289/2021) from the Universidad Politécnica de Madrid. Work performed by HM was financed by PID2020-120364GA- I00 funded by MCIN/AEI/ 10.13039/501100011033. ALG was recipient of a “María Zambrano” postdoctoral fellowship/ European Union NextGenerationEU/PRTR. We thank Gero Steinberg and Sreedhar Kilaru (Exeter University, UK) for providing us with 3D7-GFP strain and Sara González Bodí (BSU, CBGP) for providing us support with the bioinformatic analysis. We thank Cyrille Saintenac, Kostya Kanyuka and Jean-Benoit Morel for providing us with wheat seeds.

## Author contributions

DR, CCL, PK, AM and ASV designed the study. CCL, DR, PK, GL, LM, FDS and SLC performed the lab work. DR, CCL, MJM and PK analysed the data and conducted statistical analyses. AL and HM performed HPAEC-PAD analysis. CL performed the synteny plot and the population analysis. BAM contributed to the selection and functional characterization of *Zt*GH45. DR, CCL and ASV wrote the manuscript. AM, HM, AL, FV, SLC, BAM, LM, FDS, MJM and CL reviewed and edited the manuscript.

## Data availability

We declare that the raw data is in the additional files.

## Supporting Information

**Supporting Information Fig. S1 | Maps of plasmids used to overexpress *ZtGH45*. a**, Map of pRF- HU2E_ZT3D7_G10118 plasmid used for inserting *in locus* the overexpression cassette of *ZtGH45*. **b**, Map of pCCL1_ZT3D7_G10118 plasmid used for inserting ectopically the overexpression cassette of *ZtGH45*. pCCL1 was obtained by inserting the *Aspergillus nidulans* glyceraldehyde-3-phosphate dehydrogenase (*gpdA*) promoter upstream of the *TEF* terminator into the pLM1 plasmid (Meile et al. 2020) using the In-Fusion HD Cloning Kit (Takara Bio, Japan).

**Supporting Information Fig. S2 | Characterization of β-glucan oligosaccharides released during *Zymoseptoria tritici*-wheat infection.** Chromatogram and peak mass spectra of wheat leaf extracts at 10 days post infection (dpi) with *Z. tritici* using hydrophilic interaction liquid chromatography coupled to electrospray ionisation-mass spectrometry (HILIC-ESI-MS). Base peak intensity (BPI) chromatogram shown in Fig. 1a (**a**) and mass spectra of each number-labelled peak (**b**-**h**) are displayed. Degree of polymerization (DP) of putative glucose (hexose) oligosaccharides is indicated. β-ᴅ-cellobiosyl-1,3-β-ᴅ-glucose (MLG43) and cellotriose (Cello3) peaks were identified based on external standardisation.

**Supporting Information Fig. S3 | Expression pattern of *Zymoseptoria tritici* genes encoding for CAZymes with putative β-1,4-glucanase activity (EC 3.2.1.4).** Expression levels (Reads per kilobase of transcript per million reads mapped; RPKM) are shown for the strain 3D7 grown under axenic conditions (yeast extract sucrose broth (YSB) and minimal medium (MM)); and during wheat infection at 7, 12, 14 and 28 days post infection (dpi). Data were obtained from previously published RNA-seq studies (NCBI SRA accessions: SRP152081 and SRP077418).

**Supporting Information Fig. S4 | *Zymoseptoria tritici Zt*GH45 harbours a conserved catalytic domain. a**, Multiple sequence alignment using Clustal Omega of *Zt*GH45 from *Z. tritici* 3D7 SMQ54963.1 (*Zt*GH45), *Cryptopygus antarcticus* ACV50414.1 (CaGH45), *Humicola grisea* BAA74956.1 (HgGH45), *Humicola insolens* CAB42307.1 (HiGH45), *Melanocarpus albomyces* CAD56665.1 (MaGH45), *Neurospora crassa* CAD70529.1 (NcGH45) and *Thermothielavioides terrestris* AEO64667.1 (TtGH45). Signal peptides predicted by SignalP 6.0 are highlighted in red, and GH45 and cellulose-binding domains predicted by SMART in grey and green, respectively. Asterisk (*) mark conserved residues, colon (:) conservation among groups of strongly similar chemical properties and period (.) conservation among groups of weakly similar chemical properties. **b**, Cartoon representation of *Zt*GH45 structure model obtained using Alphafold2 (pLDDT: 92.7; ColabFold v1.5.5). Cysteines (yellow), residues that show variability within the *Z. tritici* Swiss population (green), residues described to mediate cellobiose-binding (cyan), and residues in the catalytic site (red), according to the *Nc*GH45 homologue protein from *N. crassa,* are shown using stick representation.

Supporting Information Fig. S5 | Early expression of *ZtGH45* impaired virulence and reproduction of *Zymoseptoria tritici* in a wheat genotype-independent manner. a, *ZtGH45* relative expression in the 3D7 wild-type strain and in the *in locus ZtGH45*-overexpression line (*ZtGH45_OE*), in which the *ZtGH45* native promoter was replaced by the constitutive promoter *gdpA* (*in locus* mutant) grown under axenic conditions. *ZtTFC1* was used as a reference gene. Bars indicate the standard error of the mean (n=3 biological replicates). *P* value according to two-tailed *t*-test between *ZtGH45_OE* line and 3D7 strain is displayed in the plot. **b**-**d**, Box plots representing the pycnidia density (pycnidia/cm^2^ of leaf) (**b;d**) and the percentage of leaf area covered by lesions (**c**) produced by 3D7 strain (control, yellow bar) and *ZtGH45_OE* lines (blue bars) on the wheat cultivar Titlis at 13 days post infection (dpi) (**b**) and on the wheat cultivar Drifter at 14 dpi (**c-d**). The data shown in **b**, correspond to the experiment shown in main Fig. 1h. *P* values according to one-way ANOVA followed by Dunnett test between plants infected with *ZtGH45_OE* and 3D7 lines are displayed in the plots.

**Supporting Information Fig. S6 | Overexpression of *ZtGH45* does not affect growth and stress tolerance. a**, Growth performance of *Zymoseptoria tritici* 3D7 strain and *in locus ZtGH45_OE* line under different stress conditions. 3 μL suspensions of 10^6^ spores/mL were grown on yeast malt sucrose agar (YMS), and YMS supplemented with 1 M sorbitol, 0.5 M NaCl, 1 mM H2O2, or 200 ng/µL Calcofluor white. Heat stress tolerance was assessed at 28 °C. **b**, Growth performance of *ZtGH45_OE* line on different carbon sources. Growth of 3D7 and *ZtGH45_OE* lines on Vogel’s minimal medium (MM) and MM supplemented with 5 g/L sucrose, 5 g/L fructose, 5 g/L carboxymethylcellulose sodium salt (CMC), or 5 g/L barley β-glucan. Images were taken 6 days after incubation at 18 °C.

**Supporting Information Fig. S7 | The catalytic site of *Zt*GH45 is required to induce resistance in wheat. a**, Relative expression of *ZtGH45* in *Zymoseptoria tritici* mutant lines that ectopically overexpress the wild-type version of *ZtGH45* (*ZtGH45_OE-WT*; blue bars) or *ZtGH45* with the mutagenized catalytic site (*ZtGH45_OE*^D37A/D145A^; red bars). Lines were grown under axenic conditions (yeast peptone dextrose, YPD). *ZtTFC1* was used as a reference gene and bars indicate the standard error of the mean (n=3 biological replicates). *P* values according to one-way ANOVA followed by Dunnett test between mutant lines and 3D7- GFP strain are displayed in the plot. **b**, Box plots representing pycnidia density (pycnidia/cm^2^ of leaf) produced on leaves of the wheat cultivar Titlis infected with 3D7-GFP strain (control, yellow bar) and independent mutant lines of *ZtGH45_OE-WT* (blue bars) or *ZtGH45_OE*^D37A/D145A^ (red bars) at 13 dpi. *P* values according to one-way ANOVA followed by Dunnett test between plants infected with mutant lines and 3D7-GFP strain are displayed in the plot.

**Supporting Information Fig. S8 | Lines of *Zymoseptoria tritici* that overexpress *ZtGH45* degrade β-glucan polysaccharides.** *Z. tritici* wild-type (3D7) and *ZtGH45-*overexpression (*ZtGH45_OE*) lines grown in Vogel’s minimal medium supplemented with 0.5% (w/v) fructose and with either 0.5% (w/v) barley mixed-linked glucan polysaccharide (β-1,3/1,4-glucan) (**a**; **c**) or 0.5% (w/v) carboxymethylcellulose sodium salt (CMC) (**b**). Pure endo-1,4-β-D-glucanase (cellulase) and water treatment were used as positive and negative controls, respectively. Oligosaccharides were quantified after 96 hours (**a**-**b**). Results from 3 independent *ZtGH45_OE* mutant lines are shown. *P* values according to one-way ANOVA followed by Dunnett test between mock, *ZtGH45_OE* lines or cellulase treatment and 3D7 strain are displayed in the plots. Bars represent the mean of four biological replicates and the error bars represent the standard error of the mean Values below detection limit are expressed as not detected (nd). Results of line 3 are shown in main Fig. 2a,b. **c**, β-1,3/1,4- glucan tetramer (MLG443), pentamer (MLG4443) and putative β-glucan oligosaccharides of high degree of polymerization (HDP) released by *Z. tritici* wild-type (3D7) and *ZtGH45_OE* lines after 96 hours of growth in β-glucan-supplemented media. The oligosaccharides were analysed by high-performance anion-exchange chromatography with pulsed amperometric detection (HPAEC-PAD). *P* values according to two-tailed *t*-test between *ZtGH45_OE* and 3D7 strain are displayed in the plot. Bars represent the mean of two biological replicates and the error bars represent the standard error of the mean. **d,** An additional replicate of the results presented in Fig. 2e. Virulence, estimated as the percentage of leaf area covered by lesions, produced by *Z. tritici* strain 3D7 at 14 days post inoculation in wheat Titlis plants pre-treated with 0.1% (v/v) UEP-100 and 0.01% (v/v) Tween 20 (Mock; grey bar), 0.5 mM β-ᴅ-cellobiosyl-1,3-β-ᴅ-glucose (MLG43, green bar) or 0.5 mM cellotriose (Cello3; red bar) for 24 hours before infection. *P* values according to Kruskal-Wallis test followed by Dunn test between the oligosaccharide treatments and the mock control are displayed in the plot.

Supporting Information Table S1. ZtGH45 annotations in different *Zymoseptoria tritici* strains. Supporting Information Table S2. Primers used in this work.

Supporting Information Table S3. Summary of the wheat and *Zymoseptoria tritici* transcriptomic dataset.

Supporting Information Table S4. Fold change (log2) expression levels of wheat genes upon MLG43, or Cello3 (3h) treatment compared to mock or upon infection with *Zymoseptoria tritici* 3D7 or *ZtGH45_OE* infected plants compared to mock treated plants (6 dpi)

Supporting Information Table S5. Counts and CPMs of wheat genes upon MLG43, or Cello3 (3h) treatment or upon infection with *Zymoseptoria tritici* 3D7 or *ZtGH45_OE* lines (6 dpi)

Supporting Information Table S6. Gene ontology (GO) analysis of up-regulated wheat genes in response to MLG43.

Supporting Information Table S7. Up-regulated wheat genes in response to MLG43. Genes potentially involved in defence are highlighted in green.

Supporting Information Table S8. Wheat genes specifically up-regulated in *ZtGH45_OE*-infected plants and not up-regulated in 3D7-infected plants. Fold change (Log2) of the expression in plants infected with *ZtGH45_OE* compared to the control are shown.

Supporting Information Table S9. Gene ontology (GO) analysis of wheat genes specifically up-regulated in *ZtGH45_OE*-infected plants and not up-regulated in 3D7-infected plants.

Supporting Information Table S10. Counts and normalized values of the expression of *Zymoseptoria tritici* genes in the wildtype and *ZtGH45_OE* lines.

Supporting Information Table S11. Differentially expressed *Zymoseptoria tritici* genes in *ZtGH45_OE*-infected plants with respect to 3D7-infected plants. Fold changes of ZtGH45_OE compared to 3D7 at 6 days post infection (dpi) are indicated (Log2).

## References

1. Alassimone J, Praz C, Lorrain C, De Francesco A, Carrasco-López C, Faino L, Shen Z, Meile L, Sanchez Vallet A. 2024. The *Zymoseptoria tritici* avirulence factor AvrStb6 accumulates in hyphae close to stomata and triggers a wheat defense response hindering fungal penetration. Molecular Plant-Microbe Interactions 37: 432–444.

2. Aziz A, Gauthier A, Bézier A, Poinssot B, Joubert JM, Pugin A, Heyraud A, Baillieul F. 2007. Elicitor and resistance-inducing activities of beta-1,4 cellodextrins in grapevine, comparison with beta-1,3 glucans and alpha-1,4 oligogalacturonides. Journal of experimental botany 58: 1463–1472.

3. Bacete L, Mélida H, Miedes E, Molina A. 2018. Plant cell wall-mediated immunity: cell wall changes trigger disease resistance responses. The Plant Journal 93: 614–636.

4. Barghahn S, Arnal G, Jain N, Petutschnig E, Brumer H, Lipka V. 2021. Mixed linkage β-1,3/1,4-glucan oligosaccharides induce defense responses in *Hordeum vulgare* and *Arabidopsis thaliana*. Frontiers in Plant Science 12: 682439.

5. Battache M, Lebrun M-H, Sakai K, Soudière O, Cambon F, Langin T, Saintenac C. 2022. Blocked at the stomatal gate, a key step of wheat Stb16q-mediated resistance to *Zymoseptoria tritici*. Frontiers in Plant Science 13: 1–14.

6. Bernasconi A, Lorrain C, Flury P, Alassimone J, McDonald BA, Sánchez-Vallet A. 2023. Virulent strains of Zymoseptoria tritici suppress the host immune response and facilitate the success of avirulent strains in mixed infections. PLOS Pathogens 19: e1011767.

7. Bigeard J, Colcombet J, Hirt H. 2015. Signaling mechanisms in pattern-triggered immunity (PTI). Molecular plant 8: 521–539.

8. Boutrot F, Zipfel C. 2017. Function, discovery, and exploitation of plant pattern recognition receptors for broad-spectrum disease resistance. Annual review of phytopathology 55: 257–286.

9. Bradley EL, Ökmen B, Doehlemann G, Henrissat B, Bradshaw RE, Mesarich CH. 2022. Secreted glycoside hydrolase proteins as effectors and invasion patterns of plant-associated fungi and oomycetes. Frontiers in Plant Science 13: 853106.

10. Brandt B, Brodsky DE, Xue S, Negi J, Iba K, Kangasjärvi J, Ghassemian M, Stephan AB, Hu H, Schroeder JI. 2012. Reconstitution of abscisic acid activation of SLAC1 anion channel by CPK6 and OST1 kinases and branched ABI1 PP2C phosphatase action. Proceedings of the National Academy of Sciences of the United States of America 109: 10593–8.

11. Bray NL, Pimentel H, Melsted P, Pachter L. 2016. Near-optimal probabilistic RNA-seq quantification. Nature Biotechnology 34: 525–527.

12. Brunner PC, Torriani SFF, Croll D, Stukenbrock EH, McDonald BA. 2013. Coevolution and life cycle specialization of plant cell wall degrading enzymes in a hemibiotrophic pathogen. Molecular biology and evolution 30: 1337–1347.

13. Burton RA, Fincher GB. 2014. Evolution and development of cell walls in cereal grains. Frontiers in Plant Scienc 5: 456–471.

14. Carpita NC, McCann M. 2000. The cell wall. In: Buchanan BB, Wilhelm G, Jones RL, eds. Biochemistry & Molecular Biology of Plants. Rockville, Illinois: American Society of Plant Physiologists, 52–108.

15. Cingolani P, Platts A, Wang LL, Coon M, Nguyen T, Wang L, Land SJ, Lu X, Ruden DM. 2012. A program for annotating and predicting the effects of single nucleotide polymorphisms, SnpEff. Fly 6: 80–92.

16. Claverie J, Balacey S, Lemaître-Guillier C, Brulé D, Chiltz A, Granet L, Noirot E, Daire X, Darblade B, Héloir MC, et al. 2018. The cell wall-derived xyloglucan is a new DAMP triggering plant immunity in vitis vinifera and *Arabidopsis thaliana*. Frontiers in Plant Science 9.

17. Costantini S, Benedetti M, Pontiggia D, Giovannoni M, Cervone F, Mattei B, De Lorenzo G. 2023. Berberine bridge enzyme–like oxidases of cellodextrins and mixed-linked β-glucans control seed coat formation. Plant Physiology 194: 296–313.

18. Dai Y-S, Liu D, Guo W, Liu Z-X, Zhang X, Shi L-L, Zhou D-M, Wang L-N, Kang K, Wang F-Z, et al. 2023. Poaceae-specific b-1,3;1,4-D-glucans link jasmonate signalling to OsLecRK1-mediated defence response during rice-brown planthopper interactions. Plant Biotechnology Journal 21: 1286–1300.

19. Deller S, Hammond-Kosack KE, Rudd JJ. 2011. The complex interactions between host immunity and non- biotrophic fungal pathogens of wheat leaves. Journal of Plant Physiology 168: 63–71.

20. Dobin A, Davis CA, Schlesinger F, Drenkow J, Zaleski C, Jha S, Batut P, Chaisson M, Gingeras TR. 2013. STAR: Ultrafast universal RNA-seq aligner. Bioinformatics 29: 15–21.

21. Drula E, Garron ML, Dogan S, Lombard V, Henrissat B, Terrapon N. 2022. The carbohydrate-active enzyme database: Functions and literature. Nucleic Acids Research 50: D571–D577.

22. Duncan KE, Howard RJ. 2000. Cytological analysis of wheat infection by the leaf blotch pathogen *Mycosphaerella graminicola*. Mycological Research 104: 1074–1082.

23. Esquerré-Tugayé MT, Boudart G, Dumas B. 2000. Cell wall degrading enzymes, inhibitory proteins, and oligosaccharides participate in the molecular dialogue between plants and pathogens. Plant Physiology and Biochemistry 38: 157–163.

24. Fernández-Calvo P, López G, Martín-Dacal M, Aitouguinane M, Carrasco-López C, González-Bodí S, Bacete L, Mélida H, Sánchez-Vallet A, Molina A. 2024. Leucine rich repeat-malectin receptor kinases IGP1/CORK1, IGP3 and IGP4 are required for arabidopsis immune responses triggered by β-1,4-D-Xylo-oligosaccharides from plant cell walls. The Cell Surface 11: 100124.

25. Förster S, Schmidt LK, Kopic E, Anschütz U, Huang S, Schlücking K, Köster P, Waadt R, Larrieu A, Batistič O, et al. 2019. Wounding-induced stomatal closure requires jasmonate-mediated activation of GORK K + channels by a Ca 2+ sensor-kinase CBL1-CIPK5 complex. Developmental Cell 48: 87–99.e6.

26. Francisco CS, Ma X, Zwyssig MM, McDonald BA, Palma-Guerrero J. 2019. Morphological changes in response to environmental stresses in the fungal plant pathogen *Zymoseptoria tritici*. Scientific Reports 9: 9642.

27. Frandsen RJN, Andersson JA, Kristensen MB, Giese H. 2008. Efficient four fragment cloning for the construction of vectors for targeted gene replacement in filamentous fungi. BMC Molecular Biology 9: 1– 11.

28. Fry SC, Nesselrode BHWA, Miller JG, Mewburn BR. 2008. Mixed-linkage (1→3,1→4)-β- D -glucan is a major hemicellulose of *Equisetum* (horsetail) cell walls. New Phytologist 179: 104–115.

29. Gámez-Arjona FM, Vitale S, Voxeur A, Dora S, Müller S, Sancho-Andrés G, Montesinos JC, Di Pietro A, Sánchez-Rodríguez C. 2022. Impairment of the cellulose degradation machinery enhances Fusarium oxysporum virulence but limits its reproductive fitness. Science Advances 8: 9734.

30. Gao J, Huang JW, Li Q, Liu W, Ko TP, Zheng Y, Xiao X, Kuo CJ, Chen CC, Guo RT. 2017. Characterization and crystal structure of a thermostable glycoside hydrolase family 45 1,4-β-endoglucanase from *Thielavia terrestris*. Enzyme and Microbial Technology 99: 32–37.

31. Geiger D, Scherzer S, Mumm P, Marten I, Ache P, Matschi S, Liese A, Wellmann C, Al-Rasheid KAS, Grill E, et al. 2010. Guard cell anion channel SLAC1 is regulated by CDPK protein kinases with distinct Ca2+ affinities. Proceedings of the National Academy of Sciences of the United States of America 107: 8023–8028.

32. Gust AA, Pruitt R, Nürnberger T. 2017. Sensing danger: key to activating plant immunity. Trends in Plant Science 22: 779–791.

33. Hahn MG, Darvill AG, Albersheim P. 1981. Host-pathogen Interactions: XIX. The endogenous elicitor, a fragment of a plant cell wall polysaccharide that elicits phytoalexin accumulation in soybeans. Plant Physiology 68: 1169.

34. Haueisen J, Stukenbrock EH. 2016. Life cycle specialization of filamentous pathogens — colonization and reproduction in plant tissues. Current Opinion in Microbiology 32: 31–37.

35. Heckman KL, Pease LR. 2007. Gene splicing and mutagenesis by PCR-driven overlap extension. Nature Protocols 2007 2:4 2: 924–932.

36. Hsu PK, Dubeaux G, Takahashi Y, Schroeder JI. 2021. Signaling mechanisms in abscisic acid-mediated stomatal closure. The Plant Journal 105: 307–321.

37. Kadowaki MAS, Polikarpov I. 2019. Structural insights into the hydrolysis pattern and molecular dynamics simulations of GH45 subfamily a endoglucanase from *Neurospora crassa* OR74A. Biochimie 165: 275–284.

38. Kema GHJ, Yu D, Rijkenberg FHJ, Shaw MW, Baayen RP. 1996. Histology of the pathogenesis of *Mycosphaerella graminicola* in wheat. Phytopathology 86: 777–786.

39. Keon J, Antoniw J, Carzaniga R, Deller S, Ward JL, Baker JM, Beale MH, Hammond-Kosack K, Rudd JJ. 2007. Transcriptional adaptation of *Mycosphaerella graminicola* to programmed cell death (PCD) of its susceptible wheat host. Molecular Plant-Microbe Interactions 20: 178–193.

40. Kong D, Hu HC, Okuma E, Lee Y, Lee HS, Munemasa S, Cho D, Ju C, Pedoeim L, Rodriguez B, et al. 2016. L- Met activates Arabidopsis GLR Ca2+ channels upstream of ROS production and regulates stomatal movement. Cell Reports 17: 2553–2561.

41. Krishnan P, Meile L, Plissonneau C, Ma X, Hartmann FE, Croll D, McDonald BA, Sánchez-Vallet A. 2018. Transposable element insertions shape gene regulation and melanin production in a fungal pathogen of wheat. BMC Biology 16: 78.

42. Kwak JM, Mori IC, Pei ZM, Leonhard N, Angel Torres M, Dangl JL, Bloom RE, Bodde S, Jones JDG, Schroeder JI. 2003. NADPH oxidase *AtrbohD* and *AtrbohF* genes function in ROS-dependent ABA signaling in Arabidopsis. The EMBO Journal 22: 2623–2633.

43. Labavitch JM, Ray PM. 1978. Structure of hemicellulosic polysaccharides of *Avena sativa* coleoptile cell walls. Phytochemistry 17: 933–937.

44. Lapalu N, Lamothe L, Petit Y, Genissel A, Delude C, Feurtey A, Abraham LN, Smith D, King R, Renwick A, et al. 2023. Improved gene annotation of the fungal wheat pathogen *Zymoseptoria tritici* based on combined Iso-Seq and RNA-Seq evidence. bioRxiv 2023.04.26.

45. Lechner M, Findeiß S, Steiner L, Marz M, Stadler PF, Prohaska SJ. 2011. Proteinortho: Detection of (Co-)orthologs in large-scale analysis. BMC Bioinformatics 12: 1–9.

46. Liao Y, Smyth GK, Shi W. 2014. featureCounts: An efficient general purpose program for assigning sequence reads to genomic features. Bioinformatics 30: 923–930.

47. Linde CC, Zhan J, McDonald BA. 2007. Population structure of *Mycosphaerella graminicola*: from lesions to continents. Phytopathology 92: 946–955.

48. Liu Y, Maierhofer T, Rybak K, Sklenar J, Breakspear A, Johnston MG, Fliegmann J, Huang S, Roelfsema MRG, Felix G, et al. 2019. Anion channel SLAH3 is a regulatory target of chitin receptor-associated kinase PBL27 in microbial stomatal closure. eLife 8: 1–23.

49. Lorrain C, Feurtey A, Ller MM, Haueisen J, Stukenbrock E. 2021. Dynamics of transposable elements in recently diverged fungal pathogens: lineage-specific transposable element content and efficiency of genome defenses. G3 Genes|Genomes|Genetics 11.

50. Lorrain C, Zurich E, Feurtey A, Alassimone J, Mcdonald B. 2024. A novel genome-wide association approach reveals wheat pathogen genes involved in host specialization. Research Square: 10.21203/rs.3.rs-4486034/v1].

51. Love MI, Huber W, Anders S. 2014. Moderated estimation of fold change and dispersion for RNA-seq data with DESeq2. Genome Biology 15: 1–21.

52. Martín-Dacal M, Fernández-Calvo P, Jiménez-Sandoval P, López G, Garrido-Arandía M, Rebaque D, del Hierro I, Berlanga DJ, Torres MÁ, Kumar V, et al. 2023. Arabidopsis immune responses triggered by cellulose- and mixed-linked glucan-derived oligosaccharides require a group of leucine-rich repeat malectin receptor kinases. The Plant Journal 113: 833–850.

53. Meile L, Peter J, Puccetti G, Alassimone J, McDonald BA, Sánchez-Vallet A. 2020. Chromatin dynamics contribute to the spatiotemporal expression pattern of virulence genes in a fungal plant pathogen. mBio 11: 1–18.

54. Mélida H, Bacete L, Ruprecht C, Rebaque D, del Hierro I, López G, Brunner F, Pfrengle F, Molina A. 2020. Arabinoxylan-Oligosaccharides Act as Damage Associated Molecular Patterns in Plants Regulating Disease Resistance. Frontiers in Plant Science 11: 557476.

55. Mélida H, Sopeña-Torres S, Bacete L, Garrido-Arandia M, Jordá L, López G, Muñoz-Barrios A, Pacios LF, Molina A. 2018. Non-branched β-1,3-glucan oligosaccharides trigger immune responses in Arabidopsis. The Plant Journal 93: 34–49.

56. Molina A, Jordá L, Torres MÁ, Martín-Dacal M, Berlanga DJ, Fernández-Calvo P, Gómez-Rubio E, Martín- Santamaría S. 2024a. Plant cell wall-mediated disease resistance: Current understanding and future perspectives. Molecular Plant 17: 699–724.

57. Molina A, Sánchez-Vallet A, Jordá L, Carrasco-López C, Rodríguez-Herva JJ, López-Solanilla E. 2024b. Plant cell walls: source of carbohydrate-based signals in plant-pathogen interactions. Current Opinion in Plant Biology 82: 102630.

58. Palma-Guerrero J, Ma X, Torriani SFF, Zala M, Francisco CS, Hartmann FE, Croll D, McDonald BA. 2017. Comparative transcriptome analyses in *Zymoseptoria tritici* reveal significant differences in gene expression among strains during plant infection. Molecular Plant-Microbe Interactions 30: 231–244.

59. Plissonneau C, Hartmann FE, Croll D. 2018. Pangenome analyses of the wheat pathogen *Zymoseptoria tritici* reveal the structural basis of a highly plastic eukaryotic genome. BMC Biology 16: 1–16.

60. Pring S, Kato H, Imano S, Camagna M, Tanaka A, Kimoto H, Chen P, Shrotri A, Kobayashi H, Fukuoka A, et al. 2023. Induction of plant disease resistance by mixed oligosaccharide elicitors prepared from plant cell wall and crustacean shells. Physiologia Plantarum 175: e14052.

61. Rebaque D, del Hierro I, López G, Bacete L, Vilaplana F, Dallabernardina P, Pfrengle F, Jordá L, Sánchez- Vallet A, Pérez R, et al. 2021. Cell wall-derived mixed-linked β-1,3/1,4-glucans trigger immune responses and disease resistance in plants. The Plant Journal 106: 601–615.

62. Rebaque D, López G, Sanz Y, Vilaplana F, Brunner F, Mélida H, Molina A. 2023. Subcritical water extraction of *Equisetum arvense* biomass withdraws cell wall fractions that trigger plant immune responses and disease resistance. Plant Molecular Biology 113: 401–414.

63. Sánchez-Vallet A, Mcdonald MC, Solomon PS, Mcdonald BA. 2015. Is *Zymoseptoria tritici* a hemibiotroph? Fungal Genetics and Biology 79: 29–32.

64. Scherzer S, Maierhofer T, Al-Rasheid KAS, Geiger D, Hedrich R. 2012. Multiple calcium-dependent kinases modulate ABA-activated guard cell anion channels. Molecular Plant 5: 1409–1412.

65. Schneider CA, Rasband WS, Eliceiri KW. 2012. NIH Image to ImageJ: 25 years of image analysis. Nature Methods 9: 671–675.

66. Sørensen I, Pettolino FA, Wilson SM, Doblin MS, Johansen B, Bacic A, Willats WGT. 2008. Mixed-linkage (1→ 3),(1 → 4)-β-D-glucan is not unique to the Poales and is an abundant component of Equisetum arvense cell walls. Plant Journal 54: 510–521.

67. Steinberg G. 2015. Cell biology of *Zymoseptoria tritici*: Pathogen cell organization and wheat infection. Fungal Genetics and Biology 79: 17–23.

68. Stewart EL, Hagerty CH, Mikaberidze A, Mundt CC, Zhong Z, McDonald BA. 2016. An improved method for measuring quantitative resistance to the wheat pathogen *Zymoseptoria tritici* using high-throughput automated image analysis. Phytopathology 106: 782–788.

69. Suarez-Fernandez M, Álvarez-Aragón R, Pastor-Mediavilla A, Maestre-Guillén A, del Olmo I, De Francesco A, Meile L, Sánchez-Vallet A. 2023. Sas3-mediated histone acetylation regulates effector gene activation in a fungal plant pathogen (A Di Pietro, Ed.). mBio 14: e0138623.

70. Vogel HJ. 1956. A convenient growth medium for *Neurospora crassa* (Medium N). Microbial Genetics Bulletin 13: 42–47.

71. Voxeur A, Habrylo O, Guénin S, Miart F, Soulié M-C, Rihouey C, Pau-Roblot C, Domon J-M, Gutierrez L, Pelloux J. 2019. Oligogalacturonide production upon *Arabidopsis thaliana*–*Botrytis cinerea* interaction. Proceedings of the National Academy of Sciences 116: 19743–19752.

72. Wickham H. 2016. Data Analysis. In: Springer-Verlag, ed. ggplot2: Elegant graphic design for data analysis. New York: Springer, Cham, 189–201.

73. Yang C, Liu R, Pang J, Ren B, Zhou H, Wang G, Wang E, Liu J. 2021. Poaceae-specific cell wall-derived oligosaccharides activate plant immunity via OsCERK1 during *Magnaporthe oryzae* infection in rice. Nature Communications 12: 1–13.

74. Zang H, Xie S, Zhu B, Yang X, Gu C, Hu B, Gao T, Chen Y, Gao X. 2019. Mannan oligosaccharides trigger multiple defence responses in rice and tobacco as a novel danger-associated molecular pattern. Molecular Plant Pathology 20: 1079.

75. Zoghlami A, Paës G. 2019. Lignocellulosic biomass: understanding recalcitrance and predicting hydrolysis. Frontiers in Chemistry 7: 874.

76. Zwiers LH, De Waard MA. 2001. Efficient *Agrobacterium tumefaciens*-mediated gene disruption in the phytopathogen *Mycosphaerella graminicola*. Current genetics 39: 388–393.

