## Supplementary Figures for "*Zymoseptoria tritici* stealth infection is facilitated by stage-specific down-regulation of a β-glucanase"

**Fig. S1 Maps of plasmids used to overexpress *ZtGH45*. a**, Map of pRF-HU2E_ZT3D7_G10118 plasmid used for inserting *in locus* the overexpression cassette of *ZtGH45*. **b**, Map of pCCL1_ZT3D7_G10118 plasmid used for inserting ectopically the overexpression cassette of *ZtGH45*. pCCL1 was obtained by inserting the *Aspergillus nidulans* glyceraldehyde-3-phosphate dehydrogenase (*gpdA*) promoter upstream of the *TEF* terminator into the pLM1 plasmid (Meile et al. 2020) using the In-Fusion HD Cloning Kit (Takara Bio, Japan).

**a**

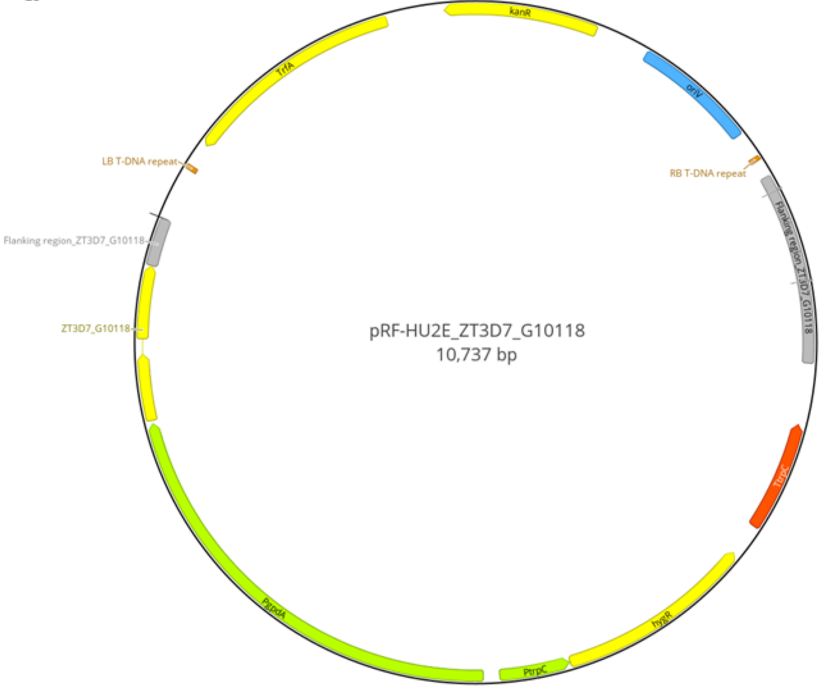
kanR
TrfA
oriV
LB T-DNA repeat
RB T-DNA repeat
Flanking region_ZT3D7_G10118
Flanking region_ZT3D7_G10118
ZT3D7_G10118
pRF-HU2E_ZT3D7_G10118
10,737 bp
TtrpC
hygR
PtrpC
PgpdA

**b**

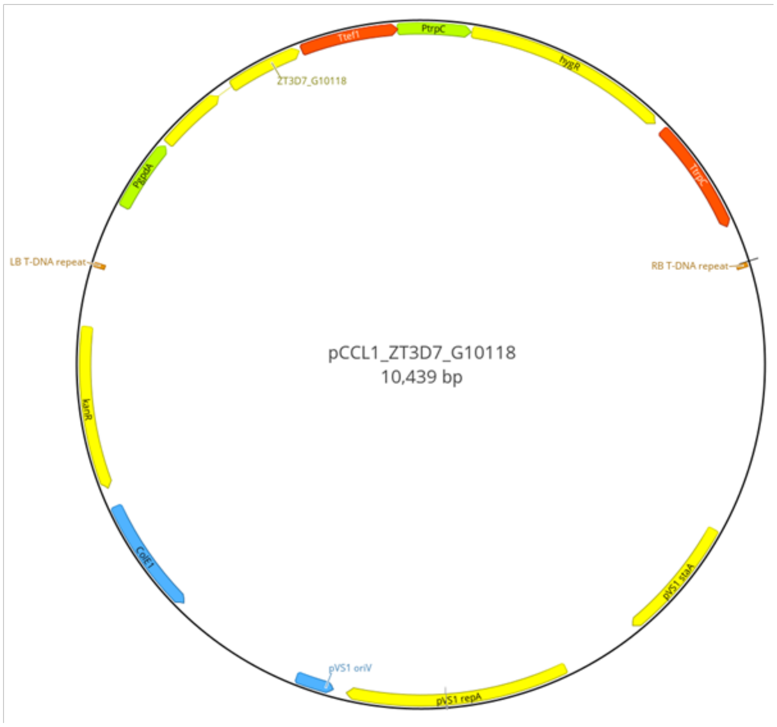
Ttef1
PtrpC
hygR
ZT3D7_G10118
PgpdA
TtrpC
LB T-DNA repeat
RB T-DNA repeat
pCCL1_ZT3D7_G10118
10,439 bp
kanR
ColE1
pVS1 staA
pVS1 oriV
pVS1 repA

**Fig. S2 Characterization of β-glucan oligosaccharides released during *Zymoseptoria tritici*-wheat infection.** Chromatogram and peak mass spectra of wheat leaf extracts at 10 days post infection (dpi) with *Z. tritici* using hydrophilic interaction liquid chromatography coupled to electrospray ionisation-mass spectrometry (HILIC-ESI-MS). Base peak intensity (BPI) chromatogram shown in Fig. 1a (**a**) and mass spectra of each number-labelled peak (**b**-**h**) are displayed. Degree of polymerization (DP) of putative glucose (hexose) oligosaccharides is indicated. β-ᴅ-cellobiosyl-1,3-β-ᴅ-glucose (MLG43) and cellotriose (Cello3) peaks were identified based on external standardisation.

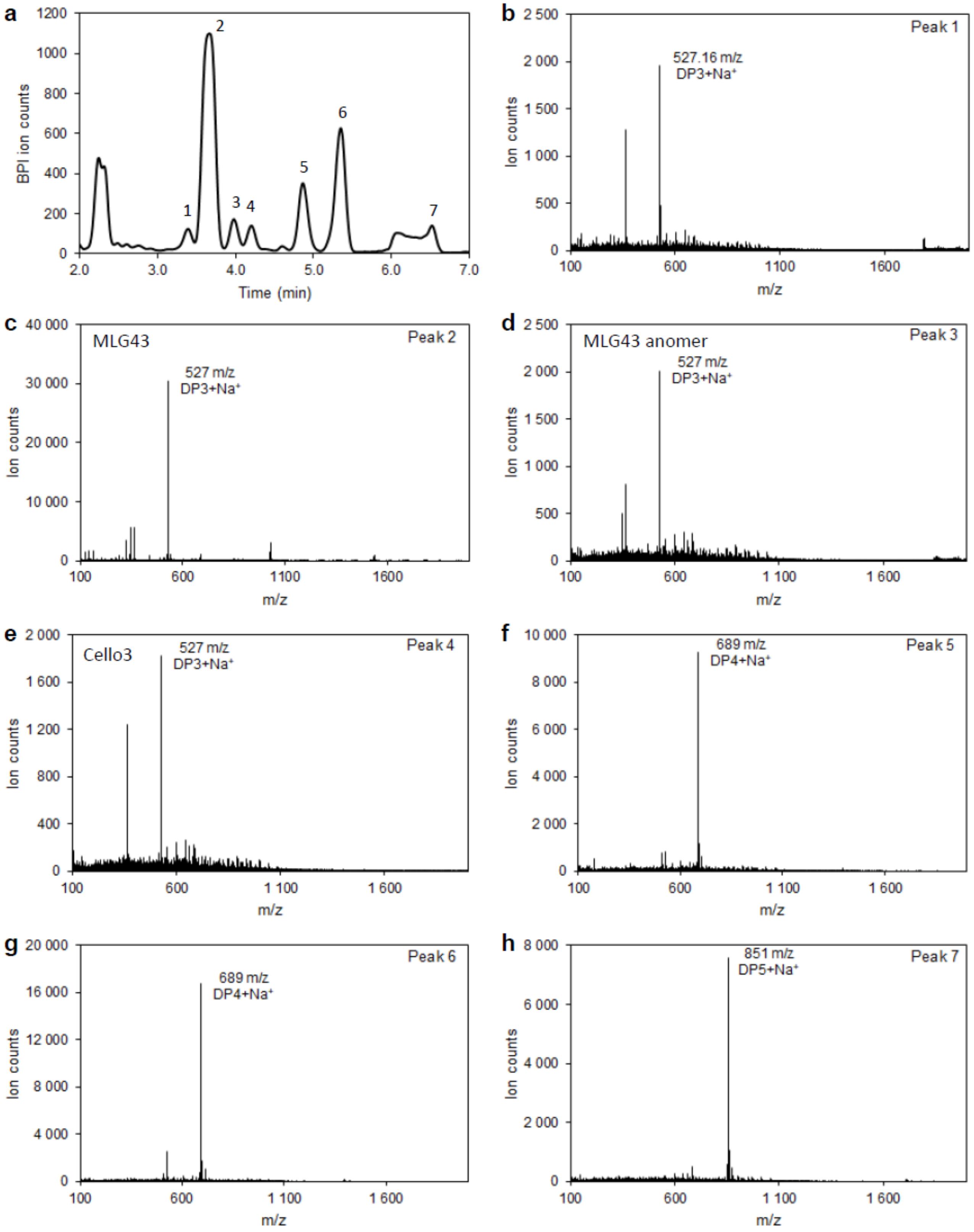

a
BPI ion counts
Time (min)
1
2
3
4
5
6
7
b
Peak 1
527.16 m/z
DP3+Na+
Ion counts
m/z
c
MLG43
Peak 2
527 m/z
DP3+Na+
Ion counts
m/z
d
MLG43 anomer
Peak 3
527 m/z
DP3+Na+
Ion counts
m/z
e
Cello3
Peak 4
527 m/z
DP3+Na+
Ion counts
m/z
f
Peak 5
689 m/z
DP4+Na+
Ion counts
m/z
g
Peak 6
689 m/z
DP4+Na+
Ion counts
m/z
h
Peak 7
851 m/z
DP5+Na+
Ion counts
m/z

**Fig. S3 Expression pattern of *Zymoseptoria tritici* genes encoding for CAZymes with putative β-1,4-glucanase activity (EC 3.2.1.4)**. Expression levels (Reads per kilobase of transcript per million reads mapped; RPKM) are shown for the strain 3D7 grown under axenic conditions (yeast extract sucrose broth (YSB) and minimal medium (MM)); and during wheat infection at 7, 12, 14 and 28 days post infection (dpi). Data were obtained from previously published RNA-seq studies (NCBI SRA accessions: SRP152081 and SRP077418).

Expression during the necrotrophic phase

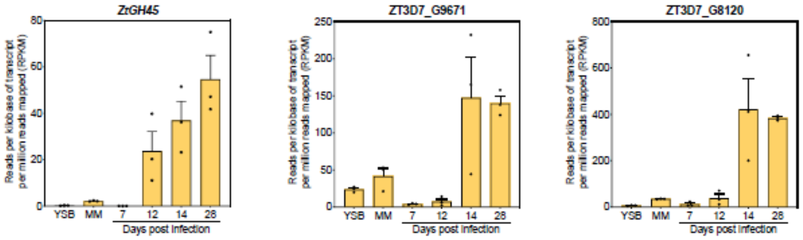


Expression throughout all the infection cycle

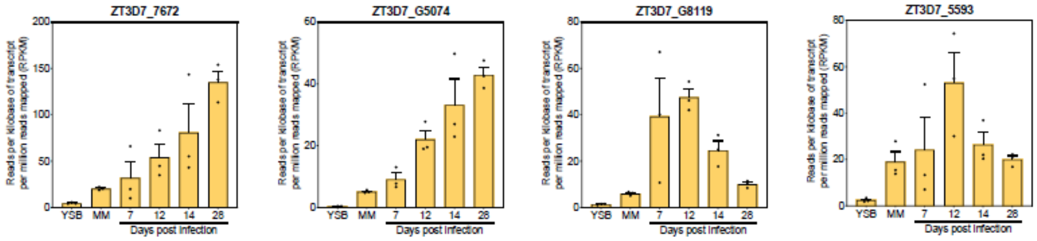


Expression at 7-12 dpi

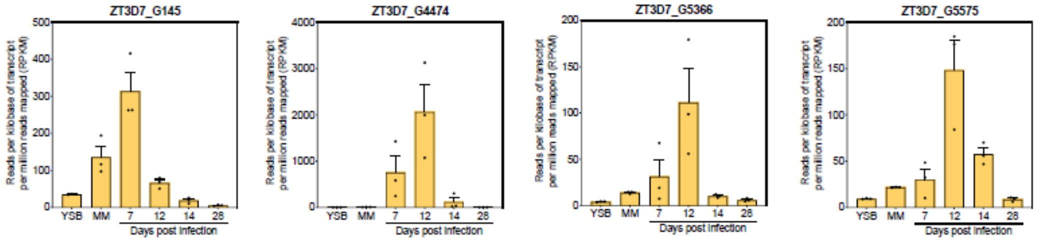


Expression under axenic conditions

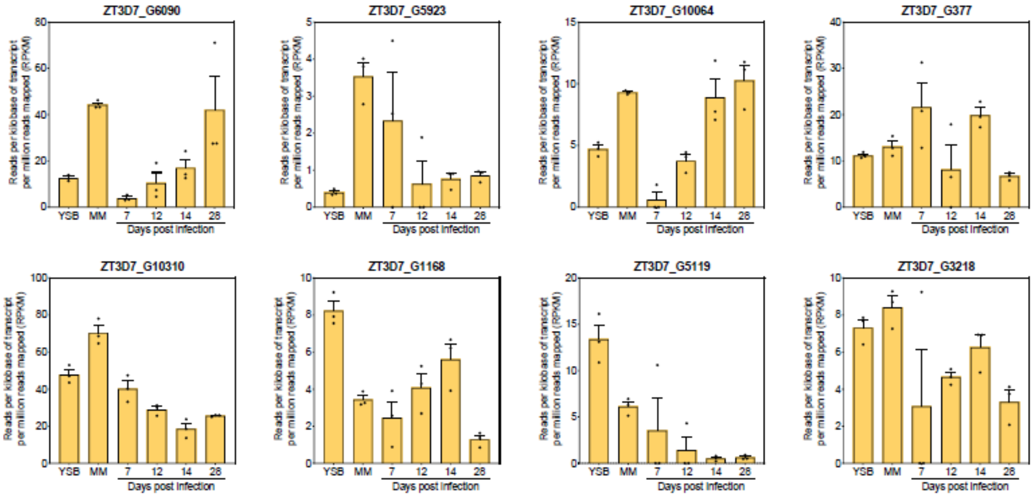

**Fig. S4 *Zymoseptoria tritici Zt*GH45 harbours a conserved catalytic domain.** Multiple sequence alignment using Clustal Omega of *Zt*GH45 from *Z. tritici* 3D7 SMQ54963.1 (*Zt*GH45), *Cryptopygus antarcticus* ACV50414.1 (CaGH45), *Humicola grisea* BAA74956.1 (HgGH45), *Humicola insolens* CAB42307.1 (HiGH45), *Melanocarpus albomyces* CAD56665.1 (MaGH45), *Neurospora crassa* CAD70529.1 (NcGH45) and *Thermothielavioides terrestris* AEO64667.1 (TtGH45). Signal peptides predicted by SignalP 6.0 are highlighted in red, and GH45 and cellulose-binding domains predicted by SMART in grey and green, respectively. Asterisk (*) mark conserved residues, colon (:) conservation among groups of strongly similar chemical properties and period (.) conservation among groups of weakly similar chemical properties. **b**, Cartoon representation of *Zt*GH45 structure model obtained using Alphafold2 (pLDDT: 92.7; ColabFold v1.5.5). Cysteines (yellow), residues that show variability within the *Z. tritici* Swiss population (green), residues described to mediate cellobiose-binding (cyan), and residues in the catalytic site (red), according to the *Nc*GH45 homologue protein from *N. crassa,* are shown using stick representation.

**a**

```
ZtGH45   MPSLSTLTLLATLTSLIQGSSLLPRQGTAGVTTRYWDCCKPSCSWPNKAAFSSPVKICGKDDNPMSDPNT   70
CaGH45   MKVFVV------LAAIV--AIANGLTSGSGVTTRYWDCCKPSCSWGGKASVTKPVRTCKANGNTTIDSNT   62
HgGH45   MRSSPLLP--SDVVAAL--PVLALA--ADGKSTRYWDCCKPSCGWAKKAPVNQPVFSCNANFQRLTDFDA   64
HiGH45   MRSSPLLP--SAVVAAL--PVLALA--ADGRSTRYWDCCKPSCGWAKKAPVNQPVFSCNANFQRITDFDA   64
MaGH45   MRSTPVLR--ALLAAAL--PLGALA--ANGQSTRYWDCCKPSCGWRGKGPVNQPVYSCDANFQRIHDFDA   64
NcGH45   MRSSTILQ--TGLVAAL--PFAVQAASGSGQSTRYWDCCKPSCSWSGKAPVNRPVLACDANNNPLSDASV   66
TtGH45   MRSTPVLR--TTLAAAL--PLVASAASGSGQSTRYWDCCKPSCAWPGKAAVSQPVYACDANFQRLSDFNV   66
         *          :.: :              * :************.*  *. .. **  *  : :   * ..

ZtGH45   VSGCD-GGPGFQCSNQRSWAKTESLAYGFAA--VNGGNEAGSCCACYKLKFTSEAAMGKTLVVQVTNIGY   137
CaGH45   QSGCN-GGSSYVCNDQQPFTQ-GNVGYGFAAASISGQPESQTCCACYEMTFTNTAISGQKMIVQVTNTGS   130
HgGH45   KSGCEPGGVAYSCADQTPWAVNDDFAFGFAATSIAGSNEAGWCCACYELTFTSGPVAGKKMVVQSTSTGG   134
HiGH45   KSGCEPGGVAYSCADQTPWAVNDDFALGFAATSIAGSNEAGWCCACYELTFTSGPVAGKKMVVQSTSTGG   134
MaGH45   VSGCE-GGPAFSCADHSPWAINDNLSYGFAATALSGQTEESWCCACYALTFTSGPVAGKTMVVQSTSTGG   133
NcGH45   KSGCD-GGSAYTCANNSPWAVNDQLSYGFAATKLSGGTESSWCCACYALTFTSGPVAGKTLVVQSTSTGG   135
TtGH45   QSGCN-GGSAYSCADQTPWAVNDNLAYGFAATSIAGGSESSWCCACYALTFTSGPVAGKTMVVQSTSTGG   135
          ***: ** .: * ::  ::   ... ****  : *  *   ***** :. **.    *:.::** *. *

ZtGH45   DVGATQFDLALPGGGVGIFPQGCRNQY-GVQNGWGSNGGYGGVSSEAECMTLPAALQDGCKFRFQWFKGS   206
CaGH45   DLNGNHFDLMIPGGGVGIFN-GCQSQWGAPSNGWGQ--RYGGISSQSECNQLPTSLRAGCNWRFGWFKNA   197
HgGH45   DLGSNHFDLNIPGGGVGIFD-GCTPQFGGLP---GQ--RYGGISSRNECDRFPDALKPGCYWRFDWFKNA   198
HiGH45   DLGSNHFDLNIPGGGVGIFD-GCTPQFGGLP---GQ--RYGGISSRNECDRFPDALKPGCYWRFDWFKNA   198
MaGH45   DLGSNHFDLNIPGGGVGLFD-GCTPQFGGLP---GA--RYGGISSRQECDSFPEPLKPGCQWRFDWFQNA   197
NcGH45   DLGSNHFDINMPGGGVGLFD-GCKRQFGGLP---GA--QYGGISSRSQCDSFPAALKPGCQWRFDWFQNA   199
TtGH45   DLGSNQFDIAMPGGGVGIFN-GCSSQFGGLP---GA--QYGGISSRDQCDSFPAPLKPGCQWRFDWFQNA   199
         *:...:**: :******:*  **  *: .       *    ***:**. :*  :*  *: ** :** **: .

ZtGH45   NNPSVMYERVACPADLVSRSGCKRDDDSRFPASP------------------------------------   240
CaGH45   DNPSMKFTQVRCPTILTQKSQCVRTPGP------------------------------------------   225
HgGH45   DNPSFSFRQVQCPAELVARTGCRRNDDGNFPAVQIPSSSTSSPVGQPTSTSTTSTSTTSSPPVQPTTPSG   268
HiGH45   DNPSFSFRQVQCPAELVARTGCRRNDDGNFPAVQIPSSSTSSPVNQPTSTSTTSTSTTSSPPVQPTTPSG   268
MaGH45   DNPSFTFERVQCPEELVARTGCRRHDDGGFAVFKAPSA--------------------------------   235
NcGH45   DNPNFTFKQVQCPSELTSRTGCKRNDDSQFPVFTPPSGGGSNPSTP-------------TTPPSSGGGSG   256
TtGH45   DNPTFTFQQVQCPAEIVARSGCKRNDDSSFPVFTPPSGGNGG-TGTPTST------APGSGQTSPGGGSG   262
         :**.. : :* **  :. :: * *   .

ZtGH45   -------------------------------------   240
CaGH45   -------------------------------------   225
HgGH45   CTAERWAQCGGNGWSGCTTCVAGSTCTKINDWYHQCL   305
HiGH45   CTAERWAQCGGNGWSGCTTCVAGSTCTKINDWYHQCL   305
MaGH45   -------------------------------------   235
NcGH45   CTADKYAQCGGSGWSGCTNCPSGSTCKTINDYYHQCA   293
TtGH45   CTSQKWAQCGGIGFSGCTTCVSGTTCQKLNDYYSQCL   299
```

**b**

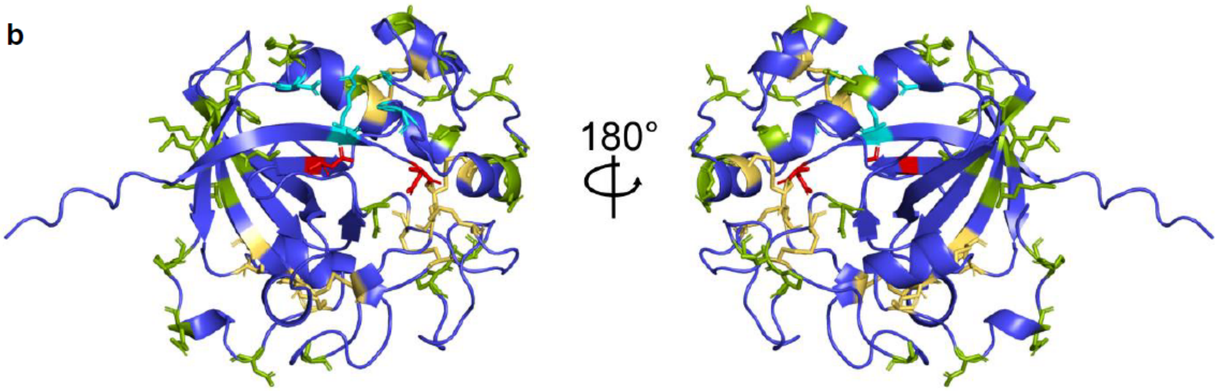

**Fig. S5 Early expression of *ZtGH45* impaired virulence and reproduction of *Zymoseptoria tritici* in a wheat genotype-independent manner. a**, *ZtGH45* relative expression in the 3D7 wild-type strain and in the *in locus ZtGH45*-overexpression line (*ZtGH45_OE*), in which the *ZtGH45* native promoter was replaced by the constitutive promoter *gdpA* (*in locus* mutant) grown under axenic conditions. *ZtTFC1* was used as a reference gene. Bars indicate the standard error of the mean (n=3 biological replicates). *P* value according to two-tailed *t*-test between *ZtGH45_OE* line and 3D7 strain is displayed in the plot. **b-d**, Box plots representing the pycnidia density (pycnidia/$cm^2$ of leaf) (**b;d**) and the percentage of leaf area covered by lesions (**c**) produced by 3D7 strain (control, yellow bar) and *ZtGH45_OE* lines (blue bars) on the wheat cultivar Titlis at 13 days post infection (dpi) (**b**) and on the wheat cultivar Drifter at 14 dpi (**c-d**). The data shown in **b**, correspond to the experiment shown in main Fig. 1h. *P* values according to one-way ANOVA followed by Dunnett test between plants infected with *ZtGH45_OE* and 3D7 lines are displayed in the plots.

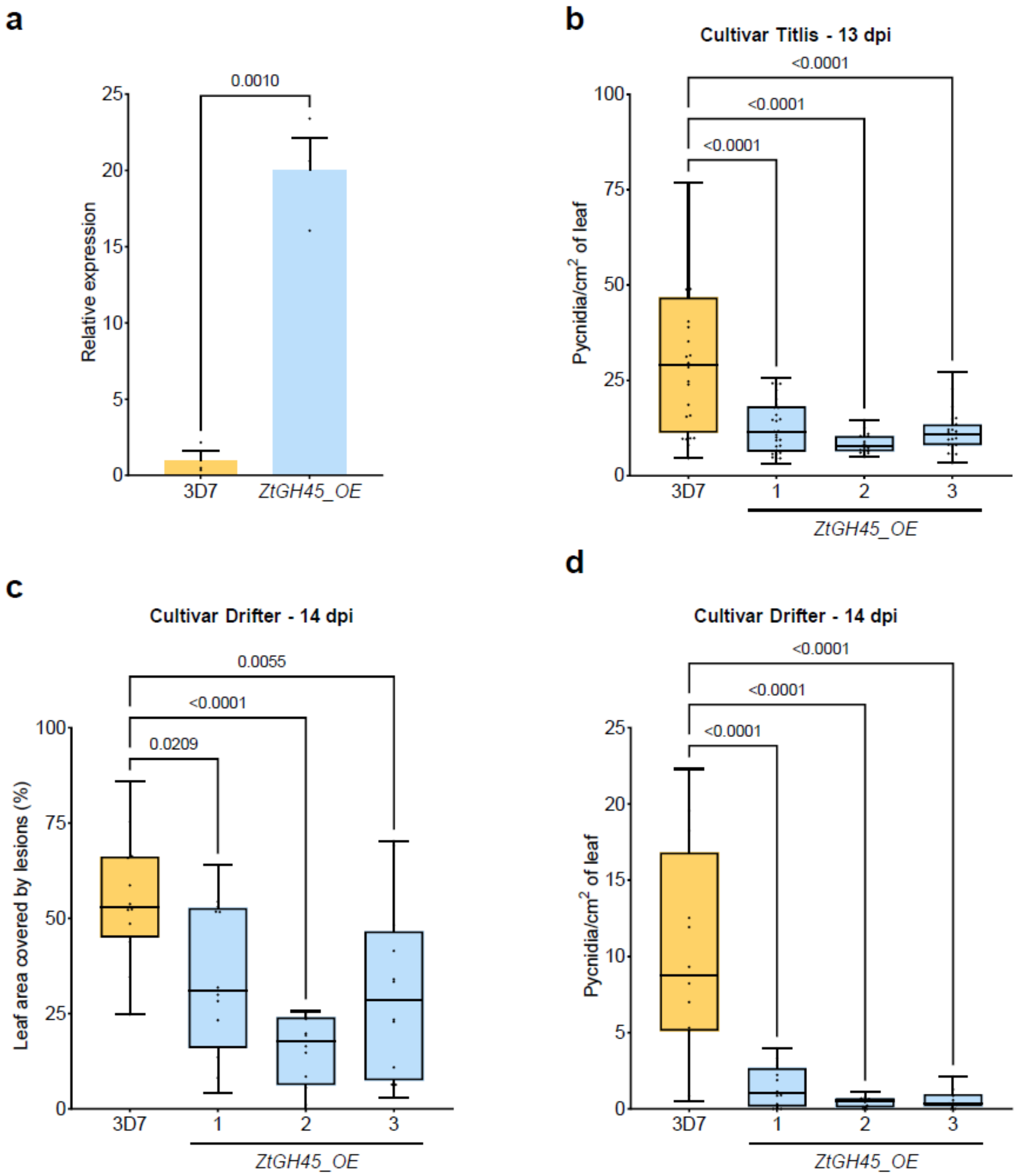
a
0.0010
25
20
15
10
5
0
Relative expression
3D7
ZtGH45_OE
b
Cultivar Titlis - 13 dpi
<0.0001
<0.0001
<0.0001
100
75
50
25
0
Pycnidia/cm2 of leaf
3D7
1
2
3
ZtGH45_OE
c
Cultivar Drifter - 14 dpi
0.0055
<0.0001
0.0209
100
75
50
25
0
Leaf area covered by lesions (%)
3D7
1
2
3
ZtGH45_OE
d
Cultivar Drifter - 14 dpi
<0.0001
<0.0001
<0.0001
25
20
15
10
5
0
Pycnidia/cm2 of leaf
3D7
1
2
3
ZtGH45_OE

**Fig. S6 Overexpression of *ZtGH45* does not affect growth and stress tolerance. a**, Growth performance of *Zymoseptoria tritici* 3D7 strain and *in locus ZtGH45_OE* line under different stress conditions. 3 μL suspensions of $10^6$ spores/mL were grown on yeast malt sucrose agar (YMS), and YMS supplemented with 1 M sorbitol, 0.5 M NaCl, 1 mM $H_2O_2$, or 200 ng/μL Calcofluor white. Heat stress tolerance was assessed at 28 °C. **b**, Growth performance of *ZtGH45_OE* line on different carbon sources. Growth of 3D7 and *ZtGH45_OE* lines on Vogel's minimal medium (MM) and MM supplemented with 5 g/L sucrose, 5 g/L fructose, 5 g/L carboxymethylcellulose sodium salt (CMC), or 5 g/L barley β-glucan. Images were taken 6 days after incubation at 18 °C.

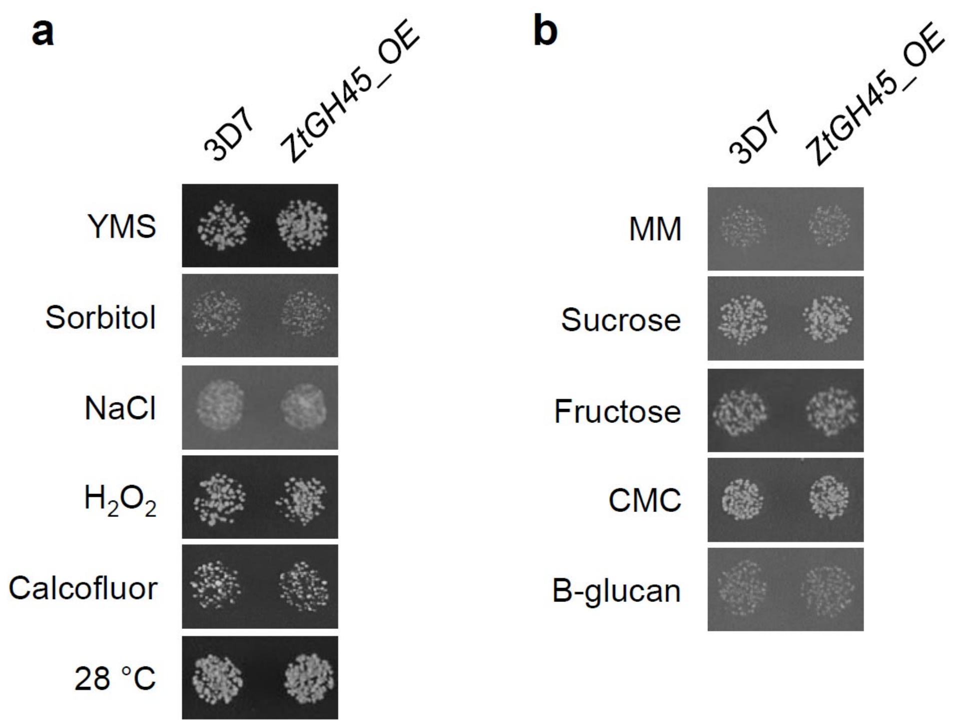

**Fig. S7 The catalytic site of *Zt*GH45 is required to induce resistance in wheat. a**, Relative expression of *ZtGH45* in *Zymoseptoria tritici* mutant lines that ectopically overexpress the wild-type version of *ZtGH45* (*ZtGH45_OE-WT*; blue bars) or *ZtGH45* with the mutagenized catalytic site (*ZtGH45_OE*$^{D37A/D145A}$; red bars). Lines were grown under axenic conditions (yeast peptone dextrose, YPD). *ZtTFC1* was used as a reference gene and bars indicate the standard error of the mean (n=3 biological replicates). *P* values according to one-way ANOVA followed by Dunnett test between mutant lines and 3D7-GFP strain are displayed in the plot. **b**, Box plots representing pycnidia density (pycnidia/cm$^2$ of leaf) produced on leaves of the wheat cultivar Titlis infected with 3D7-GFP strain (control, yellow bar) and independent mutant lines of *ZtGH45_OE-WT* (blue bars) or *ZtGH45_OE*$^{D37A/D145A}$ (red bars) at 13 dpi. *P* values according to one-way ANOVA followed by Dunnett test between plants infected with mutant lines and 3D7-GFP strain are displayed in the plot.

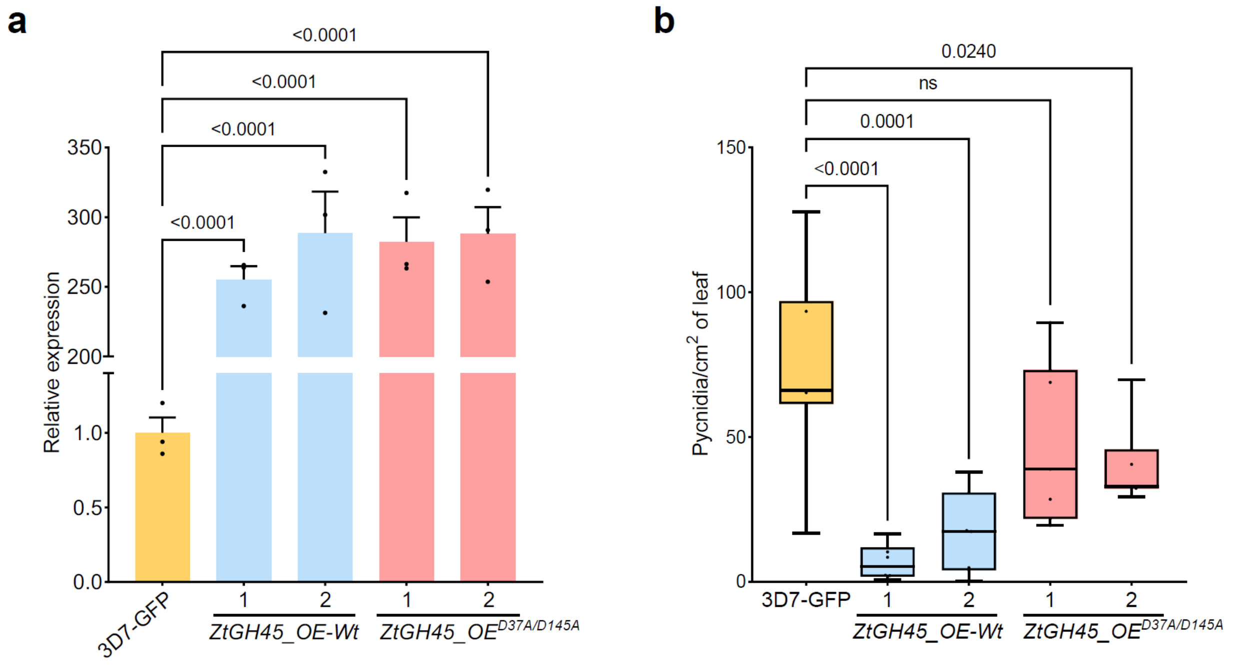

**Fig. S8 Lines of *Zymoseptoria tritici* that overexpress *ZtGH45* degrade β-glucan polysaccharides.** *Z. tritici* wild-type (3D7) and *ZtGH45*-overexpression (*ZtGH45_OE*) lines grown in Vogel's minimal medium supplemented with 0.5% (w/v) fructose and with either 0.5% (w/v) barley mixed-linked glucan polysaccharide (β-1,3/1,4-glucan) (**a**; **c**) or 0.5% (w/v) carboxymethylcellulose sodium salt (CMC) (**b**). Pure endo-1,4-β-D-glucanase (cellulase) and water treatment were used as positive and negative controls, respectively. Oligosaccharides were quantified after 96 hours (**a**-**b**). Results from 3 independent *ZtGH45_OE* mutant lines are shown. *P* values according to one-way ANOVA followed by Dunnett test between mock, *ZtGH45_OE* lines or cellulase treatment and 3D7 strain are displayed in the plots. Bars represent the mean of four biological replicates and the error bars represent the standard error of the mean Values below detection limit are expressed as not detected (nd). Results of line 3 are shown in main Fig. 2a,b. **c**, β-1,3/1,4-glucan tetramer (MLG443), pentamer (MLG4443) and putative β-glucan oligosaccharides of high degree of polymerization (HDP) released by *Z. tritici* wild-type (3D7) and *ZtGH45_OE* lines after 96 hours of growth in β-glucan-supplemented media. The oligosaccharides were analysed by high-performance anion-exchange chromatography with pulsed amperometric detection (HPAEC-PAD). *P* values according to two-tailed *t*-test between *ZtGH45_OE* and 3D7 strain are displayed in the plot. Bars represent the mean of two biological replicates and the error bars represent the standard error of the mean. **d,** An additional replicate of the results presented in Fig. 2e. Virulence, estimated as the percentage of leaf area covered by lesions, produced by *Z. tritici* strain 3D7 at 14 days post inoculation in wheat Titlis plants pre-treated with 0.1% (v/v) UEP-100 and 0.01% (v/v) Tween 20 (Mock; grey bar), 0.5 mM β-D-cellobiosyl-1,3-β-D-glucose (MLG43, green bar) or 0.5 mM cellotriose (Cello3; red bar) for 24 hours before infection. *P* values according to Kruskal-Wallis test followed by Dunn test between the oligosaccharide treatments and the mock control are displayed in the plot.

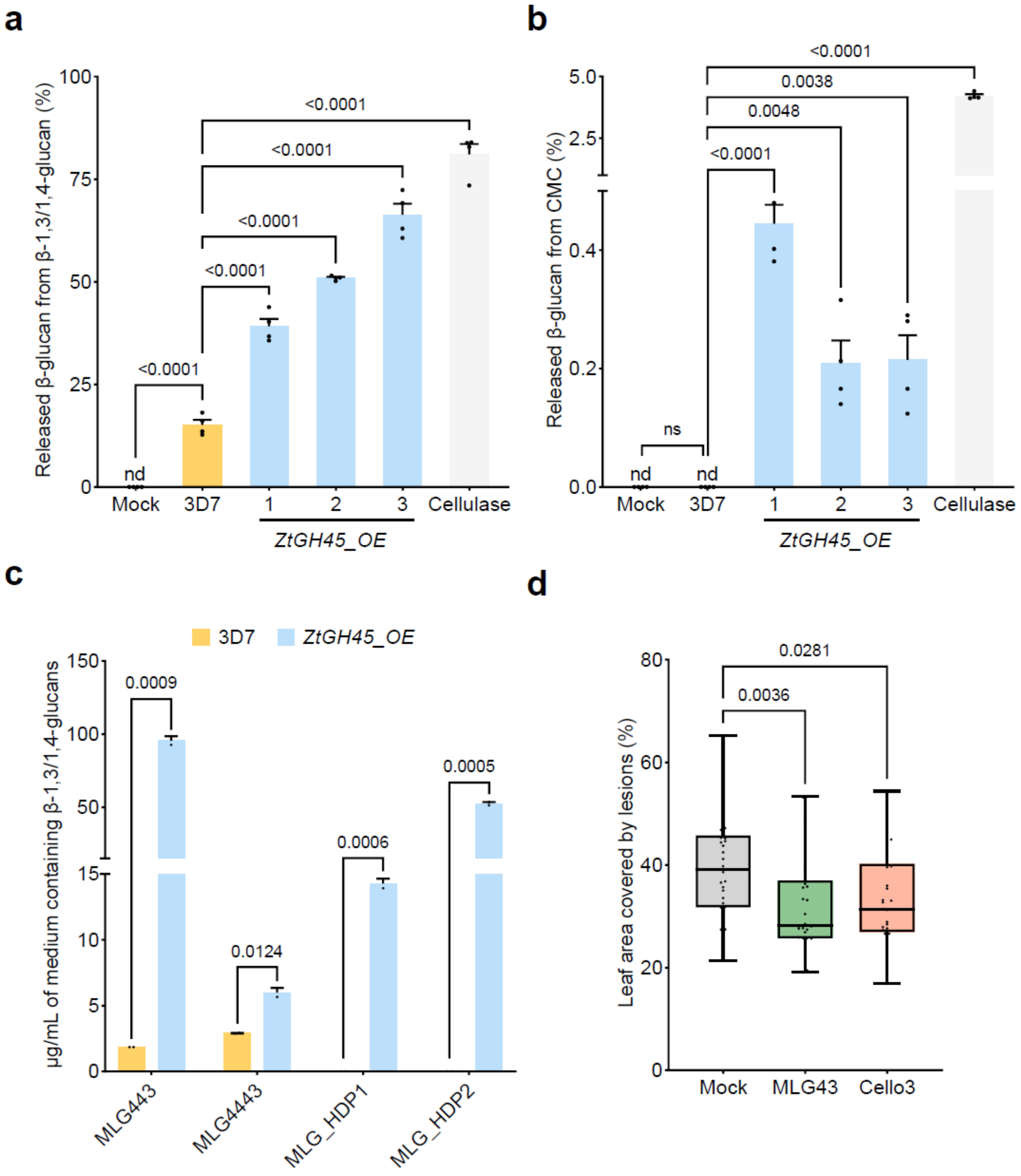
